# Polymorphism-aware models in RevBayes: Species trees, disentangling Balancing Selection and CG-biased gene conversion

**DOI:** 10.1101/2023.12.11.571102

**Authors:** Svitlana Braichenko, Rui Borges, Carolin Kosiol

## Abstract

The role of balancing selection is a long-standing evolutionary puzzle. Balancing selection is a crucial evolutionary process that maintains genetic variation (polymorphism) over extended periods of time; however, detecting it poses a significant challenge. Building upon the polymorphism-aware phylogenetic models (PoMos) framework rooted in the Moran model, we introduce PoMoBalance model. This novel approach is designed to disentangle the interplay of mutation, genetic drift, directional selection (GC-biased gene conversion), along with the previously unexplored balancing selection pressures on ultra-long timescales comparable with species divergence times by analysing multi-individual genomic and phylogenetic divergence data. Implemented in the open-source RevBayes Bayesian framework, PoMoBalance offers a versatile tool for inferring phylogenetic trees as well as quantifying various selective pressures. The novel aspect of our approach in studying balancing selection lies in PoMos’ ability to account for ancestral polymorphisms and incorporate parameters that measure frequency-dependent selection, allowing us to determine the strength of the effect and exact frequencies under selection. We implemented validation tests and assessed the model on the data simulated with SLiM and a custom Moran model simulator. Real sequence analysis of *Drosophila* populations reveals insights into the evolutionary dynamics of regions subject to frequency-dependent balancing selection, particularly in the context of sex-limited colour dimorphism in *Drosophila erecta*.

## 1 Introduction

Balancing selection (BS) represents a form of natural selection that maintains beneficial genetic diversity within populations (Bitarello *et al*., 2023). Multiple mechanisms contribute to maintaining variation, such as the heterozygote advantage or overdominance (heterozygous individuals having higher fitness), frequency-dependent selection (an individual’s fitness depends on the frequencies of other phenotypes or genotypes), antagonistic selection (in contexts like sexual conflicts or tissue-specific antagonism) and selection that changes through time or space in population. The evidence for BS is extensive, including examples from immune response such as the major histocompatibility complex (MHC) (Andrés *et al*., 2009; Spurgin and Richardson, 2010; Bitarello *et al*., 2018), pathogen resistance (Bakker *et al*., 2006), plant and fungi self-incompatibility (Lawrence, 2000; Castric and Vekemans, 2004), and sex-related genes (Charlesworth, 2004; Connallon and Clark, 2014; Mank, 2017; Kim *et al*., 2019).

BS finds its roots in the “balance hypothesis”, according to which populations exhibited high levels of diversity, with natural selection maintaining a balance among different alleles (Dobzhansky, 1955). Historically, the classical theory diminished the ubiquity of the balancing hypothesis by explaining evolution of populations through the interplay of mutations and purifying or positive selections with varying strengths. However, balancing selection remains a valuable concept for explaining the persistence of polymorphisms over extended periods. According to Bitarello *et al*. (2023), three types of balancing selection can be defined based on the acting timescales. Assuming the effective population size (*N*_*e*_ = 10^6^ (Sprengelmeyer *et al*., 2020)), generation time (10 days (Fernández-Moreno *et al*., 2007)) and the divergence times between *Drosophila erecta* and *Drosophila orena* species (3 *×* 10^6^ years (Yassin *et al*., 2016)), which are studied here, one can translate these timescales into calendar times. In this context, balancing selection can be categorized as ultra long-term (*>* 3.7 *×* 10^6^ years), long-term (*>* 10^5^ years) and recent (*<* 10^5^ years).

The heterozygote advantage stands out as one of the initially proposed mechanisms for balancing selection. The textbook example of this kind of balancing selection is found in human African populations: homozygous individuals for the abnormal version of *β*-globin gene that makes hemoglobin are susceptible to sickle-cell disease, while heterozygous individuals exhibit resistance to malaria (Laval *et al*., 2019). In this study, even though we are capable of detecting heterozygote advantage as well, we focus more on another well-known mechanism of balancing selection called negative frequency-dependent balancing selection as defined by Charlesworth and Charlesworth (2010). This mechanism is observed when the fitness of one individual depends on the frequencies of other phenotypes or genotypes in the population. Very often, negative frequency-dependent selection manifests in the maintenance of one or several rare advantageous genotypes in a population. In the context of this study, we focus on ultra long-term balancing selection (*∼* 5 million years), which leads to sexual dimorphism in female *Drosophila erecta* resulting in the maintenance of dark and light females in the populations. The dark females are presumably engaging in mimicry among the males to avoid the costs associated with repeated matings (Yassin *et al*., 2016).

The role of BS has been a subject of considerable debate over the last century (Bitarello *et al*., 2023). With the advent of new sequencing technologies, there has been a renewed interest in this phenomenon. Some models, such as those based on heterozygote advantage and sexual antagonism, have been proposed by Connallon and Clark (2014); Zeng *et al*. (2021). While these models are valuable for describing allele frequency dynamics in populations, they become impractical for inference due to the consideration of specific cases of BS that are challenging to generalise and increasing computational costs associated with expanding parameter space.

Thus, a model that is flexible enough to capture the intricate effects of BS yet simple is required for inferring frequency-dependent selection. Here, we propose a new model that incorporates BS and further integrates it into an inference approach. We build upon PoMos, a set of models developed over a decade for species tree inference (De Maio *et al*. (2013, 2015); Schrempf *et al*. (2016)). A fast implementation of the PoMo approach for species tree inference is available in IQ-TREE (Schrempf *et al*., 2019).

Recently, PoMos were extended to account for directional selection (DS) and tested on the GC-biased gene conversion (gBGC) (Borges *et al*., 2019; Borges and Kosiol, 2020; Borges *et al*., 2022a,b). This phenomenon is modelled similarly to DS, by setting relative fitnesses for *C* and *G* alleles higher than those for *A* and *T* alleles. Furthermore, in the inference setup, DS and gBGC are considered to be equivalent.

Borges and colleagues (Borges *et al*., 2022a) demonstrated that including the effect of gBGC improves the accuracy of branch length estimation employed for molecular dating. Here, in the context of BS, we integrate the modelling of gBGC as it serves as the background force. This approach provides a more realistic null model, thereby enhancing the estimation of BS on experimental data.

PoMos prove to be valuable for modelling and detecting BS, as they are rooted in polymorphisms characterized by the prolonged existence of multiple genetic variations — markers of BS (Bitarello *et al*., 2023). This phenomenon manifests in a shift in the site frequency spectrum (SFS) towards an excess of intermediate frequency variants. These are sometimes identifiable by a peak in the intermediate frequencies of the SFS that cannot be explained by the interplay between mutation, genetic drift, and directional selection, as mentioned in Charlesworth (2006); Charlesworth and Charlesworth (2010) but by BS. Consequently, these signatures are utilised by various frameworks to detect BS.

BS poses a significant challenge to detection methods due to its subtle nature, often entangled with structural variants and linkage disequilibrium (Charlesworth, 2006; Fijarczyk and Babik, 2015). Recent efforts have been made to propose universal and robust frameworks for BS detection. The software packages aimed at detecting balancing selection are summarized in Table 1. These include methods based on genome scans with multiple summary statistics and composite likelihood ratio tests (CLRT) (Andrés *et al*., 2009; DeGiorgio *et al*., 2014; Bitarello *et al*., 2018; Cheng and DeGiorgio, 2019, 2020, 2022), as well as deep-learning methods (Sheehan and Song, 2016; Isildak *et al*., 2021; Korfmann *et al*., 2023). In Table 1, we summarize approaches that are most relevant to our study; for more details, please refer to Bitarello *et al*. (2023).

**Table 1:** Comparison of PoMos with other methods for detection of DS and BS.

| Function<br>Package | Citation | Method | Multi-<br>species | Trained<br>and<br>tested<br>on human | DS | Detects<br>magnitude<br>and<br>frequencies |
| --- | --- | --- | --- | --- | --- | --- |
| BetaScan | Siewert and Voight (2017)<br>Siewert and Voight (2020) | Summary<br>Statistics | - | + | - | - |
| BALLET | DeGiorgio *et al.* (2014) | CLRT | - | + | - | - |
| NCD | Bitarello *et al.* (2018) | Summary<br>Statistics | - | - | - | - |
| MuteBaSS | Cheng and DeGiorgio (2019) | Summary<br>Statistics | + | - | - | - |
| MULLET | Cheng and DeGiorgio (2019) | CLRT | + | + | - | - |
| BaLeRMix | Cheng and DeGiorgio (2020) | CLRT | - | + | - | - |
| BaLeRMix+ | Cheng and DeGiorgio (2022) | CLRT | - | + | + | - |
| BaSe | Isildak *et al.* (2021) | Deep<br>Learning | - | + | + | - |
| PoMos<br>with selection<br>(PoMoSelect +<br>PoMoBalance) | Borges *et al.* (2019)<br>Borges *et al.* (2022b)<br>This study | Bayesian<br>Inference | + | - | + | + |

The majority of the approaches mentioned above exploit long-term BS and are therefore focused on scenarios involving single species. Two exceptions to this are MuteBaSS and MULLET (Cheng and DeGiorgio, 2019), which operate within the paradigm of ultra-long BS and accept multi-species data. Consequently, we utilise these packages for comparisons with our approach. Another aspect of the methods summarised in Table 1, is that the majority of them are trained and tested on human or great ape data. Therefore, one must exercise caution when applying them to other species. Moreover, unlike other approaches, Cheng and DeGiorgio (2022) strives to disentangle DS from BS. However, their approach requires intricate information about populations, such as recombination maps and ancestral pairwise alignment files.

By leveraging the advantages of accommodating multi-species data, applicability to most species (excluding bacteria and viruses), and incorporating mechanisms for disentangling DS from BS, our approach serves as a Bayesian inference tool. Our method not only detects selection but also quantifies its strength and frequencies, unlike most of the BS detection tools that show maximal performance at frequency equilibrium close to 0.5. Notably, NCD (Bitarello *et al*., 2018) and subsequently MuteBaSS (Cheng and DeGiorgio, 2019), which utilises modified NCD statistics, possess a mechanism to detect BS at frequency equilibrium below 0.5. However, these frequencies must be pre-defined by the user. BetaScan2 (Siewert and Voight, 2020) is also capable of detecting equilibrium frequencies, but when substitutions are specified, it is outperformed by NCD (Cheng and DeGiorgio, 2019).

Evaluating the effect of BS remains challenging, requiring more model-based approaches (Fijarczyk and Babik, 2015; Bitarello *et al*., 2023). Specifically, we require models that extend beyond heterozygote advantage, incorporating frequency-dependent selection, and integrating both balancing and directional selection. Our method addresses these challenges in a particular manner. Currently, it focuses on single genes or groups of genes; however, it holds a high potential for parallel implementation. It allows analyses across numerous individuals and populations over genomic regions including several hundred base pairs now and in the future it is poised to enable genome-wide inferences.

## 2 Materials and Methods

### 2.1 Modelling the Balancing Selection with PoMoBalance

In this paper, we introduce the PoMoBalance model (depicted in Figure 1 (A)) that can be regarded as an extension of the PoMos with DS introduced by Borges *et al*. (2019); Borges and Kosiol (2020); Borges *et al*. (2022a,b). We will refer to the latter as PoMoSelect henceforth for brevity. Both PoMoSelect and PoMoBalance are distinguished in Table 3 and belong to the family of models known as PoMos, that are continuous-time Markov chain models based on the Moran model (Moran, 1958). The Moran model is a stochastic process that simulates a virtual population of *N* haploid individuals, with the power to incorporate boundary mutations and directional selection. Together with the Wright-Fisher model, they are both boundary mutation models. These models treat mutations as perturbations from the equilibrium state of populations, while selection drives population genotypes to fixation. The frequency-dependent formulation of such models makes them attractive for inference, since it is relatively easy to implement DS and BS in them. Moran model bears similarities to the Wright-Fisher model, which counts time in the number of generations. In contrast, the Moran model is continuous-time, measuring time in the number of births (Lanchier, 2017). This characteristic makes the Moran model advantageous for phylogeny and experimental evolution approaches (Barata *et al*., 2023) that rely on a continuous-time paradigm.

**Figure 1:**
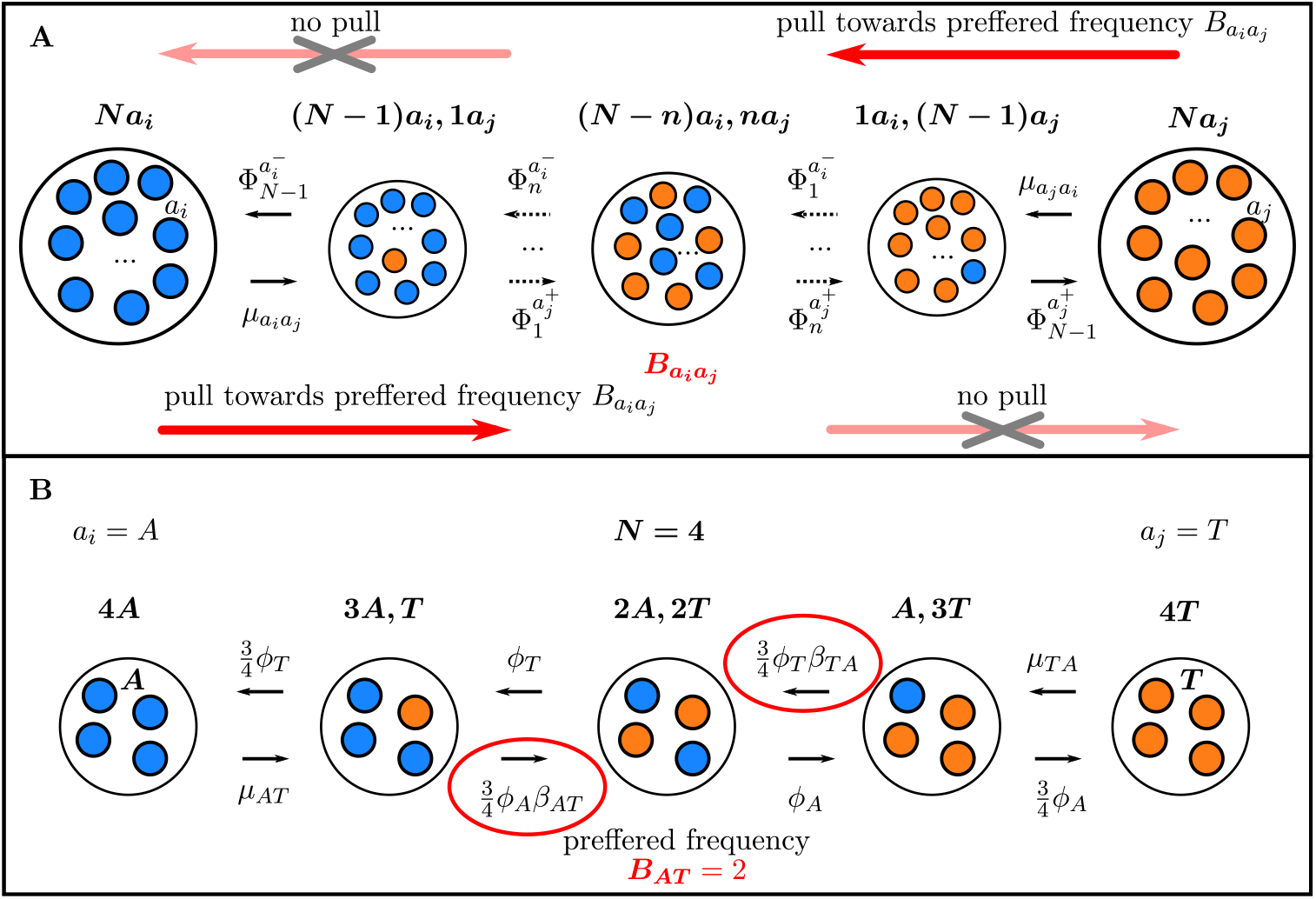
(A) PoMoBalance model, presented as a Markov chain Moran-based model. The boundary states (monomorphic) are denoted by larger circles. These states encompass *N* individuals, with the left side showcasing individuals carrying the *a*_*i*_ allele (depicted as blue circles), and the right side representing individuals with the *a*_*j*_ allele (represented by orange circles). In contrast, all the intermediate states, reflecting polymorphic conditions, are displayed using smaller circles. The transition rates from the monomorphic states are determined by mutation rates, whereas the transition rates from the polymorphic states are governed by the multiplicative fitness as indicated in Equation (1). Additionally, the multiplicative fitness encapsulates not only the DS effect but also the influence of BS, which exerts a force towards the state with the preferred allele frequency, 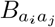, represented by dark red arrows. If the transition occurs against this preferred state, there is no such attracting force, signified by the light red crossed arrows. (B) A specific instance of the PoMoBalance model, featuring a population size of *N* = 4.

In this paper, we extend the Moran model to include balancing selection in a four-allelic system representing the four nucleotide bases. The model encompasses 4 + 6(*N* − 1) distinct states, with four monomorphic boundary states, denoting scenarios in which all individuals share the same allele. In contrast, the intermediate 6(*N* − 1) states represent polymorphisms, where some individuals possess different alleles. Here, as shown in Figure 1 (A), we consider only biallelic polymorphisms, where each state represents certain frequency *n* of alleles *a*_*i*_ (depicted as blue circles) and *N* − *n* of *a*_*j*_ (depicted as orange circles). These alleles signify four nucleotides *i, j* = *{A, C, G, T }*. The combinations of alleles, indicated as *a*_*i*_*a*_*j*_, represent the possible pairs without repetition, namely *AC, AG, AT*, *CG, CT*, or *GT* .

The model incorporates mutation rates, 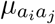 and 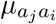 (as illustrated in Figure 1 (A)), which govern transitions from the monomorphic states, representing boundary mutations. The parameters of PoMos are defined in Table 2. Very often, the reversibility of the model is defined from certain symmetries in the parameters. In PoMoSelect the mutation rates are presented as 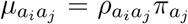 and 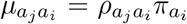, similar to Tavare (1986). Parameters 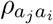 are exhange-abilities of nucleotides (Yang, 2014), that specify the relative rates of change between states *i* and *j*, and 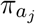 are nucleotide base frequencies, giving the equilibrium frequency at which each base occurs at all sites. If 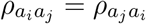 the model is reversible, otherwise, it is non-reversible.

**Table 2:** Parameters of PoMos in the four-allelic case.

| Parameter | Variable or vector | Description |
| --- | --- | --- |
| $a_{i,j}$ | $A, C, G, T$ | nucleotide bases |
| $a_i a_j$ | $AC, AG, AT, CG, CT, GT$ | pairwise combinations of nucleotide bases |
| $N$ | $N$ | effective population size |
| $\mu_{a_i a_j}$ | $\mu = (\mu_{AC}, \mu_{AG}, \mu_{AT}, \mu_{CG}, \mu_{CT}, \mu_{GT}, \mu_{CA}, \mu_{GA}, \mu_{TA}, \mu_{GC}, \mu_{TC}, \mu_{TG})$ | mutation rates |
| $\mu_{a_i a_j} = \rho_{a_i a_j} \pi_{a_j}$ | $\mu = (\mu_{AC}, \mu_{AG}, \mu_{AT}, \mu_{CG}, \mu_{CT}, \mu_{GT})$ | reversible mutation rates |
| $\pi_{a_{i,j}}$ | $\pi = (\pi_A, \pi_C, \pi_G, \pi_T)$ | nucleotide base frequencies |
| $\rho_{a_i a_j}$ | $\rho = (\rho_{AC}, \rho_{AG}, \rho_{AT}, \rho_{CG}, \rho_{CT}, \rho_{GT})$ | exchangeabilities |
| $\phi_{a_{i,j}}$ | $\phi = (\phi_A, \phi_C, \phi_G, \phi_T)$ | fitnesses |
| $\sigma_{a_{i,j}} = \phi_{a_{i,j}} - 1$ | $\sigma_{sel} = (\sigma_A, \sigma_C, \sigma_G, \sigma_T)$<br>[if $\sigma_A = \sigma_T = 0, \sigma = \sigma_C = \sigma_G$] | selection coefficients<br>[GC-bias rate] |
| $\beta_{a_i a_j}$ | $\beta = (\beta_{AC}, \beta_{AG}, \beta_{AT}, \beta_{CG}, \beta_{CT}, \beta_{GT})$ | strength of BS |
| $B_{a_i a_j} [B_{a_i a_j}/N]$ | $B = (B_{AC}, B_{AG}, B_{AT}, B_{CG}, B_{CT}, B_{GT})$ | preffered [equilibrium] frequencies of BS |

**Table 3:** PoMo functions and parameters in RevBayes.

| Function (Reference in the text) | Description | Parameters |
| --- | --- | --- |
| `fnPomoKN`<br>(non-reversible PoMoSelect or PoMoSelect) | Describes the evolution of a population with $K$ alleles and $N$ individuals subjected to mutational bias and selection. | $K, N, \mu, \phi$ |
| `fnReversiblePomoKN`<br>(reversible PoMoSelect) | Particular case of `PoMoKN` when mutations are considered reversible. | $K, N, \pi, \rho, \phi$ |
| `fnPomoBalanceKN`<br>(non-reversible PoMoBalance or PoMoBalance) | Describes the evolution of a population with $K$ alleles and $N$ individuals subjected to mutational bias, selection and balancing selection. | $K, N, \mu, \phi, \beta, B$ |
| `fnReversiblePomoBalanceKN`<br>(reversible PoMoBalance) | Particular case of `PoMoBalanceKN` when mutations are considered reversible and the preferred frequency is in the middle $B = \frac{N}{2}$. | $K, N, \pi, \rho, \phi, \beta$ |

In PoMoSelect, frequency shifts between polymorphic states are governed by genetic drift and directional selection favouring or disfavouring the reproduction of the *a*_*i*_ allele. The fitness values are represented by 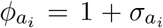, where 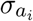 is a selection coefficient. In PoMoBalance, these frequency shifts additionally include balancing selection transition rates that are regulated by a quantity that we call multiplicative fitness, expressed by the following equation for the selected state *n* as per Figure 1 (A)

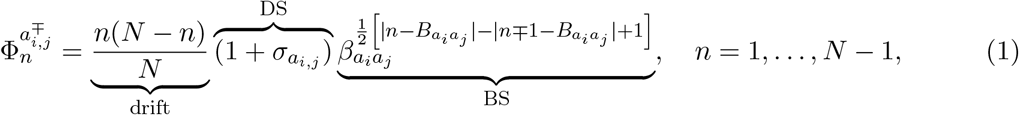

where there are three components: the first fraction corresponds to genetic drift or neutral mutations, the second multiplier represents directional selection, modelled similarly to previous PoMos. The final term in the form of a power-law function characterizes BS. This form of the BS term was derived phenomenologically from observations of various SFS derived from experimental data. It is governed by two key factors: the strength of BS, denoted as 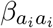 (with 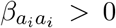), and a preferred frequency denoted as 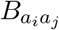. The preferred frequency, a natural number within the range 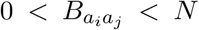, designates the position of the polymorphic peak associated with BS in the SFS. Note that if 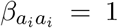 the resulting model aligns with the PoMoSelect model. We modelled BS in a frequency-dependent manner, in which the strength of balancing selection governing the frequency shifts towards a favoured frequency. The frequency equilibrium, as defined in Charlesworth and Charlesworth (2010); Bitarello *et al*. (2018, 2023); Andrés *et al*. (2009), can be determined from our model as 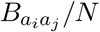 .

Reversibility criteria for PoMoBalance are different from those for the PoMoSelect model due to the higher complexity of the transition rates from the polymorphic states brought by balancing selection terms. PoMoBalance is reversible only if exhangeabilities are symmetric and the preferred frequency is in the middle of the chain 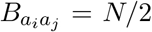, where *N* is even (for more details see Section Supplementary Material 1).

Furthermore, we always assume that both 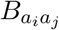 and 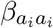 are symmetric. The strength of BS operates similarly to directional selection, but rather than favouring the fixation of alleles, it promotes the persistence of polymorphisms. In Figure 1 (A), we visualize this additional attraction towards the preferred polymorphic state with dark red arrows when 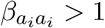. After replacing variables and simplifying the expressions with power terms, the transition rates become 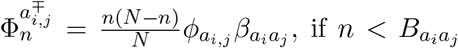, and the absence of the BS attractor is indicated with light red crossed arrows in the figure when 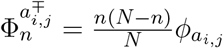, if 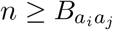. To provide a more concrete example, we represent the transition rates of a population with *N* = 4 individuals in Figure 1 (B), where the preferred frequency is *B* = 2. It is important to note that in cases where 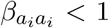, we do not model balancing selection, but instead a form of purging selection occurs that leads to the removal of polymorphisms more than expected by drift (for a detailed explanation see Section Supplementary Material 1).

In the broader context, the PoMoBalance model can be characterised through the instantaneous rate matrix denoted as *Q*, where each specific transition rate within the model corresponds to an element of this matrix

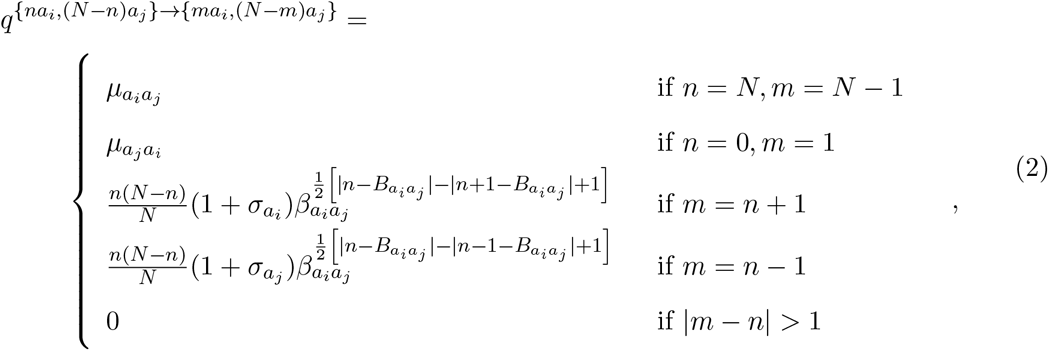

where the variables *n* and *m* represent neighbouring states as illustrated in Figure 1 (A). This matrix summarises PoMoBalance model, depicting transitions from monomorphic states regulated by mutation rates and from polymorphic states governed by Equation 1. Since PoMoBalance is Moran-based model, the allele frequency shifts exceeding one are prohibited, as specified in the final condition outlined in Equation (2). The diagonal elements of this matrix are determined such that the sum of each respective row is equal to zero.

Both the PoMoSelect and PoMoBalance models have been incorporated into a Bayesian phylogenetic inference framework RevBayes (Höhna *et al*., 2016; Hohna *et al*., 2017; Höhna *et al*., 2018; Borges *et al*., 2022b), available at https://revbayes.github.io/, employing a probabilistic graphical model representation.

### 2.2 Bayesian Inference using PoMoBalance with RevBayes

The advantage of using RevBayes for implementing PoMos is the flexibility of the use of probabilistic graphical models allowing us to combine complex models while taking advantage of communicating them with users through extensive tutorials and discussion forums. RevBayes employs a Bayesian inference based on the Markov chain Monte Carlo (MCMC) sampler and it is an open-source framework for phylogenetic inference, molecular dating, discrete morphology and ancestral state reconstruction (Höhna *et al*., 2016; Hohna *et al*., 2017; Höhna *et al*., 2018; Borges *et al*., 2022b). Our integration of PoMoBalance into RevBayes enables users to perform phylogenetic tree inference, directional selection analysis, and now, identify balancing selection. Unlike previous methods for detecting balancing selection discussed earlier, our software not only detects balancing selection but also quantifies its strength and identifies the alleles and their frequencies under selection. For detailed instructions on implementing RevBayes scripts with PoMoBalance, please refer to the PoMoBalance tutorial available at https://revbayes.github.io/tutorials/pomobalance/.

In PoMos’ data input, count files are employed, which can be generated from FASTA sequences of multiple individuals and species or VCF files with the corresponding reference using the cflib package available on GitHub at https://github.com/pomo-dev/cflib (Schrempf *et al*., 2016). Additionally, we include scripts to correct for sampling biases, which can be helpful when the number of individuals sampled from populations varies and when it differs from the PoMo population size. These biases may emerge from undersampling genetic diversity, where polymorphic sites sampled from larger populations may erroneously appear monomorphic. To address this, the binomial sampling method, as initially proposed by Schrempf *et al*. (2016), assists in smoothing out sampling biases at the tips of a phylogenetic tree.

Additionally, PoMoSelect includes a rescaling tool for adjusting inferred parameters across different population sizes. Parameters calculated in the PoMos, originally in terms of virtual population sizes, can be rescaled to represent the actual population sizes. This rescaling is achieved using the mapping method introduced by Borges *et al*. (2019) and explained in the context of PoMoBalance in Section Supplementary Material 2.

RevBayes offers several PoMo functions tailored to different inference scenarios, including `fnPomoKN, fnReversiblePomoKN, fnPomoBalanceKN and fnReversiblePomoBalanceKN`. The first two functions are discussed in detail by Borges *et al*. (2022b). The roles and input parameters for each function are summarised in Table 3. They are designed to infer data from *K* alleles, with the most common scenario involving *K* = 4, although other options (e.g., *K* = 2) are also available. Additionally, RevBayes accommodates the parameters of the PoMoBalance model outlined in Subsection 2.1 and Table 2. These include the virtual population size *N*, mutation rates *μ* represented through nucleotide base frequencies *π* and exhangeabilities *ρ* in the reversible case. Additionally, it includes a vector encompassing allele fitnesses *φ*, which, in our case, reflects gBGC as previously studied by Borges *et al*. (2019). We sometimes mention DS and gBGC interchangeably since the latter is modelled similarly to DS, with higher relative fitness for *C* and *G* alleles compared to *A* and *T* alleles. It is represented by the vector *φ* = (1, 1 + *σ*, 1 + *σ*, 1), where *σ* is a GC-bias rate. We also define two vectors for the strength and location of the balancing selection peak for each combination of alleles *β* and *B*.

For the Bayesian inferences conducted here, we employ `dnDirichlet` priors (concentration 0.25 for all alleles) on base frequencies *π* and `mvBetaSimplex` moves due to their sum-to-unity nature. For *ρ, σ* and *β* `dnExponential` priors are chosen as appropriate priors for positive real parameters with rates 10, 10 and 1, respectively, similar for all combinations of alleles. We use standard `mvScale` moves for these variables, but if they exhibit correlation, we may introduce additional moves like `mvUpDownScale, mvAVMVN, mvSlice` or `mvEllipticalSliceSamplingSimple` to mitigate the correlation. In some cases, we observed a correlation between *σ* and *β*, and incorporating the `mvAVMVN` move helped to resolve it for some chains. The preferred frequency *B* is a positive natural number within the range 0 *< B < N*, and `Uniform` priors in this range are set. The variable is rounded on each MCMC step to obtain discrete results. We introduce two moves, `mvSlide` and `mvScale`, to enhance parameter space exploration. Such a technique leads to faster convergence compared to `UniformNatural` prior and discrete variable moves. We assign different weights to each move; however, the specific values are less critical since autotuning of weights occurs during the MCMC burn-in period. Our analysis involves running both the Metropolis-Hastings MCMC sampler (`mcmc`), and where relevant, the Metropolis-coupled MCMC sampler or MC^3^ (`mcmcmc`), which includes high-temperature and cold chains to overcome local minima. Both versions normally run 4 parallel chains to ensure convergence. The number of MCMC steps required for convergence (ESS *>* 200) for different types of analyses is depicted in Figure 2.

**Figure 2:**
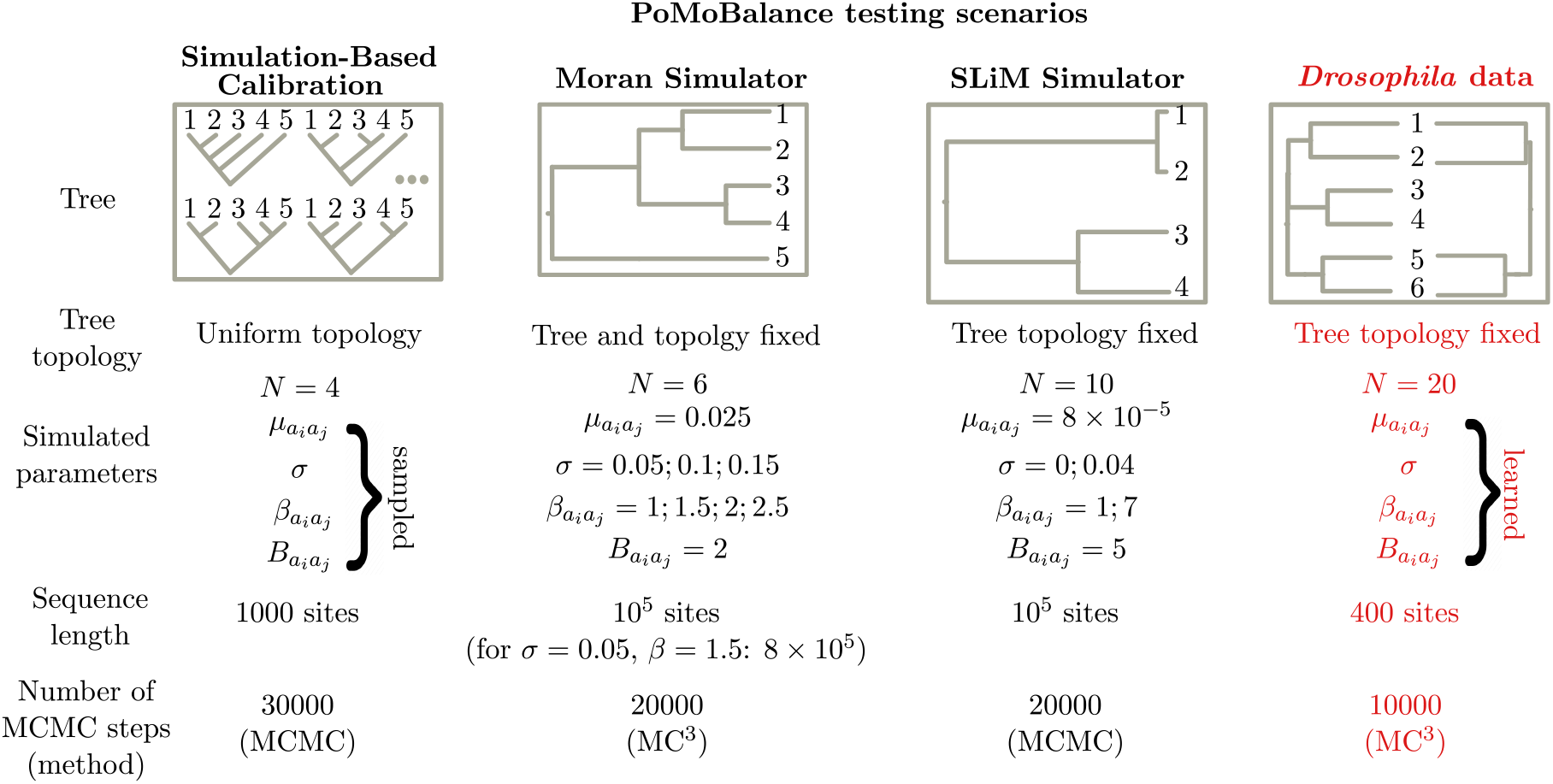
Testing scenarios for PoMoBalance include various types of trees, tree topologies, parameters of PoMos utilised in the tests, sequence lengths, and the number of MCMC steps. Simulation-based calibration involves data simulated under 1000 parameters sampled from priors, while the Moran and SLiM frameworks also rely on simulated data for several values of *σ* and *β*. Additionally, we employ experimental data extracted from various subspecies of *Drosophila*.

### 2.3 Data Simulation, Analysis and Inference

Extensive testing of PoMoBalance has been conducted across multiple scenarios, employing data simulated through different techniques. Each scenario is summarised in Figure 2.

Firstly, we conducted a built-in validation analysis within RevBayes. This analysis is based on the Simulation-Based Calibration procedure (Talts *et al*., 2020), the approach used to test the accuracy of parameter inference through the following steps:

1. Drawing 1000 random parameter values and a random five-species trees with uniform topology from the priors.
2. For each drawn parameter value simulating data sample with 1000 nucleotide sites.
3. Performing MCMC inference for each sample.
4. Calculating coverage probabilities.

Coverage probabilities (Talts *et al*., 2020) are estimated based on the observation that 90% (or any arbitrary percentage) of credible intervals obtained with MCMC should contain the simulated parameter value in 90% of the samples. SBC leverages the frequentist properties of Bayesian inference. The advantage of this approach is its ability to simultaneously test the model across various parameters and multiple five-species trees. Additionally, we calculate the scores for tree topology, measured by mean Robinson-Foulds distances (Höhna *et al*., 2018)), inferring tree topologies especially for large trees known to be a notoriously challenging task (Cavalli-Sforza and Edwards, 1967). The deliberate choice of a small virtual population size, *N* = 4, aims to test our models with minimal computational cost, as previous findings suggest that performance tends to remain consistent even with an increase in *N* (Borges et al., unpublished). We also conducted tests with *N* = 6, yielding similar performance. However, testing with higher values of *N* becomes challenging due to the increasing computational cost associated with larger values. Nonetheless, we anticipate that the performance would remain consistent.

Subsequently, a custom five-species tree (refer to Figure 2 and Figure 4 (A)) was simulated using a Moran simulator in RevBayes. In this analysis, we utilise a five-species tree, as most methods for detecting ultra long-term BS focus on testing fixed trees with four species or fewer (Cheng and DeGiorgio, 2019). Our simulations cover timescales associated with long-term or ultra-long-term balancing selection, as this example is not tied to any specific species. Here, we maintain the tree fixed to ensure better performance of the method. We recommend employing PoMoSelect for tree inference initially, as it has demonstrated better performance in inferring tree topologies (refer to Figure 3). In testing the PoMoBalance approach, our focus is primarily on inferring gBGC and balancing selection parameters. We simulate the sequences under the same model to ensure the precise recovery of parameters from data simulated under the similar models but in diverse evolutionary settings, including drift, CG-biased gene conversion, balancing selection, and a combination of balancing selection and gBGC. For most of the values, we simulated 10^5^ genomics sites, while for the intertwined scenario of weak BS 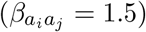 and gBGC (*σ* = 0.05), we required 8 *×* 10^5^ to achieve satisfactory convergence.

**Figure 3:**
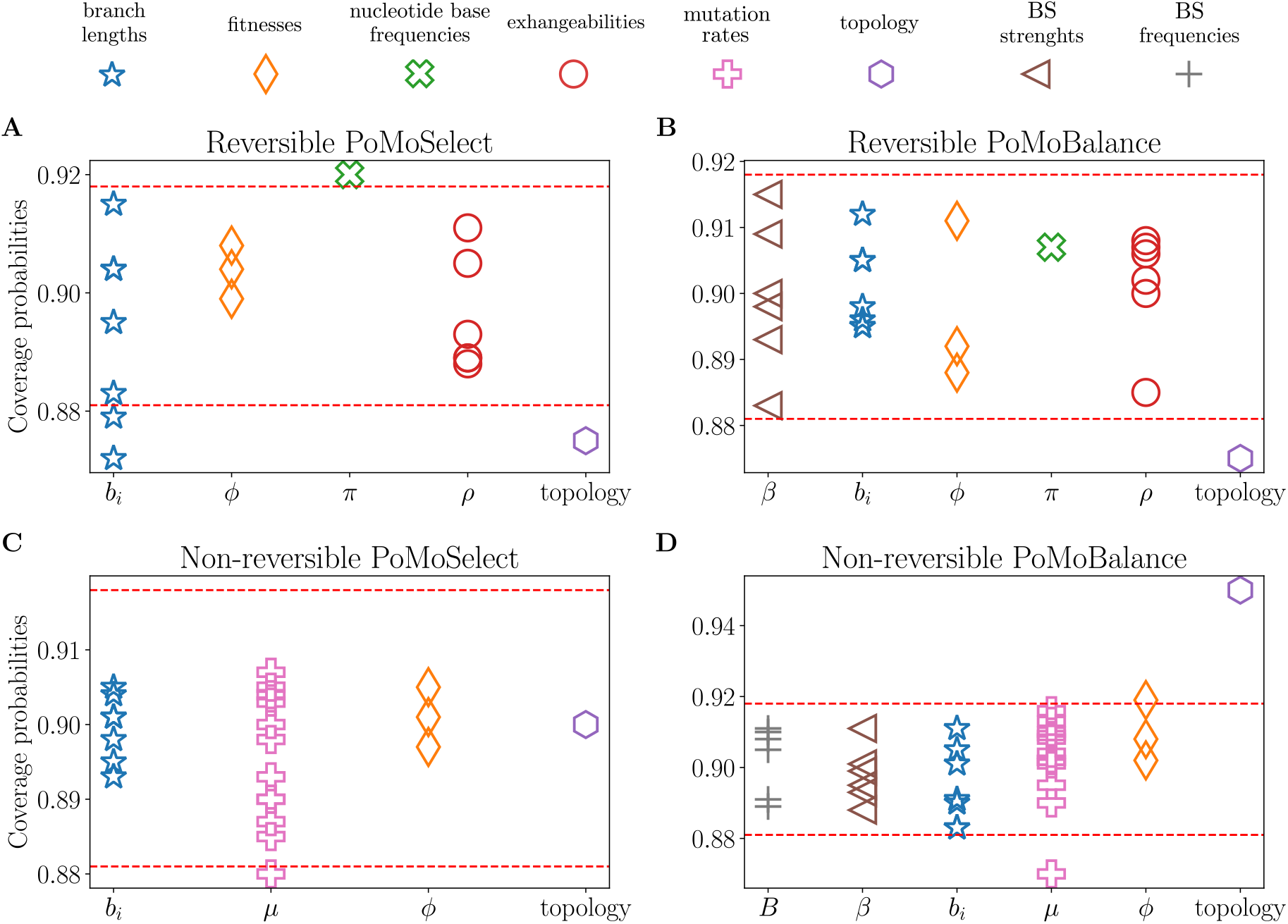
Coverage probabilities determined through validation analysis within RevBayes, employing distinct computational routines for reversible scenarios: (A) PoMoSelect and (B) PoMoBalance, as well as for non-reversible scenarios: (C) PoMoSelect and (D) PoMoBalance. The red dashed lines indicate 90% confidence intervals and fixed virtual population size for all cases was *N* = 4.

**Figure 4:**
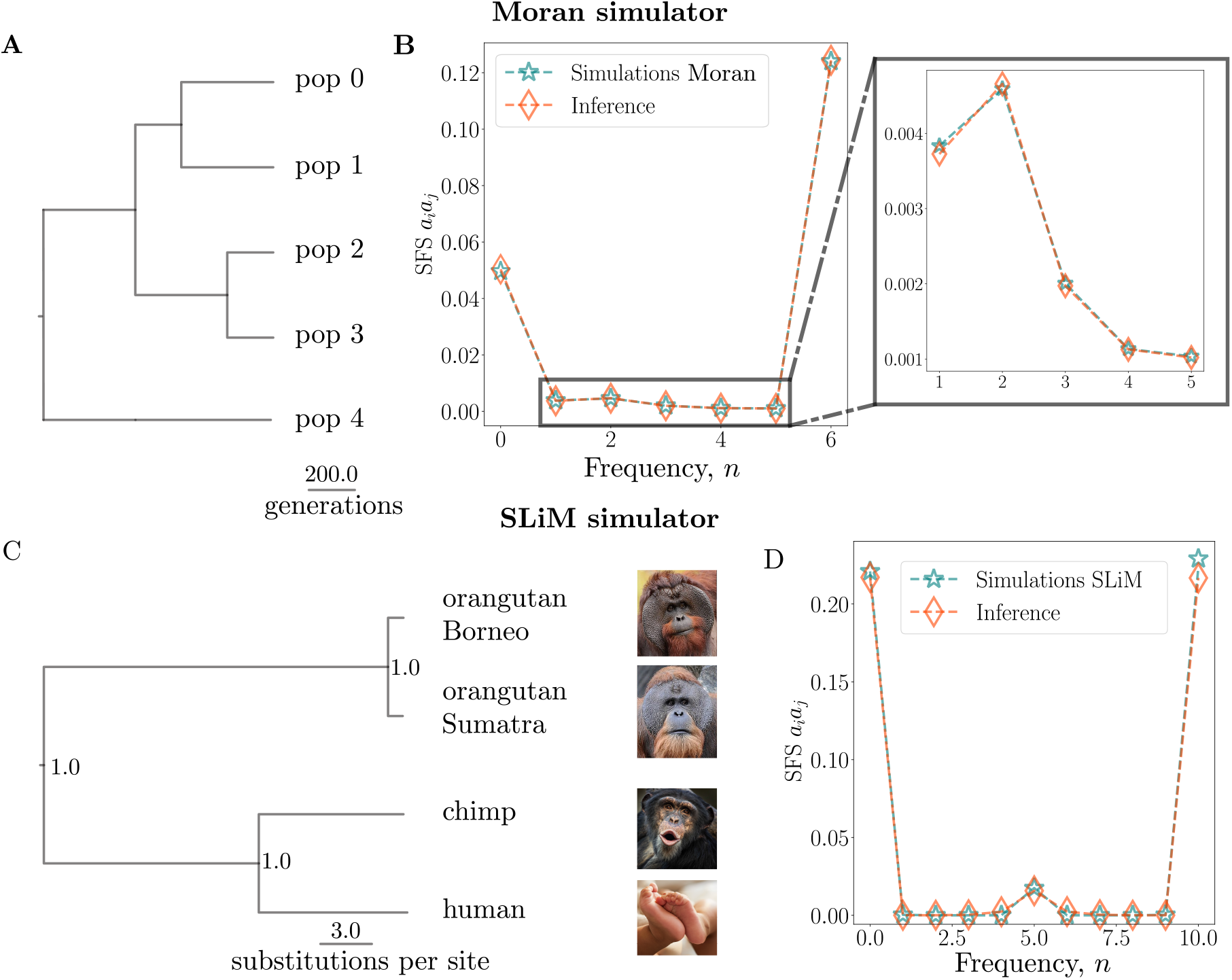
(A) Phylogenetic tree simulated using the Moran simulator within RevBayes, the branch lengths are expressed in numbers of generations; the tree remains fixed for these analyses. (B) Site-frequency spectrum of the data with balancing selection simulated using the Moran model with *N* = 6 (blue stars), with the tree from (A) exhibiting good agreement with the SFS obtained from the inference using PoMoBalance (orange diamonds); the inset magnifies the BS peak. (C) Phylogenetic tree of great apes simulated with SLiM and subsequently inferred with RevBayes, the branch lengths are expressed in the number of substitutions per site. Posterior probabilities are indicated at the nodes. Images are distributed under a Creative Commons license from Wikimedia and Microsoft. (D) Comparison of the SFS with *N* = 10, akin to (B), obtained from the simulated data with SLiM and the tree from (C). The SFS representation (*a*_*i*_*a*_*j*_) includes *AC, AG, AT*, *CG, CT* and *GT*, demonstrating similarity in all cases.

Furthermore, we assessed the performance of our package using data simulated within the evolutionary framework SLiM (Haller and Messer, 2019). In this test, we used a tree including four great ape species: orangutans from Borneo and Sumatra islands, chimpanzees, and humans (refer to Figure 2, Figure 4 (C) and Supplementary Figure S1). This tree had been previously estimated without balancing selection using PoMos by Schrempf *et al*. (2016). In this setup, we simulate ultra long-term BS and we first infer the tree with PoMoSelect. Subsequently, for the PoMoBalance analysis, we maintain the tree topology fixed and infer tree branch lengths alongside other parameters. The great ape species are of particular interest in the context of our paper as they exhibit several well-documented instances of balancing selection, such as those observed in the MHC locus (Cagan *et al*., 2016). Another classical example of heterozygote advantage is sickle-cell disease, extensively studied in humans, however, its role in other great ape species remains a subject of debate (Laval *et al*., 2019). In SLiM simulations, we implemented heterozygote advantage within the great apes tree to simulate balancing selection. Unlike the Moran simulator, SLiM simulations incorporated three regimes: drift, gBGC, and BS, excluding combination of BS and gBGC. This adjustment was necessary due to the heterozygote advantage overpowering gBGC in SLiM. Other features not explicitly considered by the Moran model but simulated in SLiM are genetic recombination and demography. Refer to Section Supplementary Material 3 for more details on SLiM simulations.

Following this, we applied PoMoBalance to real datasets exhibiting balancing selection associated with sexual dimorphism in *Drosophila erecta* females (Yassin *et al*., 2016). This case was chosen to exemplify ultra long-term negative frequency-dependent balancing selection in sexual selection, a topic of increasing interest (Croze *et al*., 2017). Please refer to Figure 2 for details of the inference and Section 7 for data availability details. Sequences were obtained for the *tan* gene in the t_MSE_ region. In addition to *Drosophila erecta* dark (7 individuals) and light (9 individuals), we extract data of multiple individuals from four closely related subspecies: *D. santomea* (10 individuals), *yakuba* (15 individuals), *melanogaster* (22 individuals) and *simulans* (18 individuals). We inferred trees in two cases: when all six subspecies were involved, and in the four-subspecies case, where we discarded *D. santomea* and *yakuba* due to poor quality of sequences. We performed the sequence alignment using MAFFT software (Rozewicki *et al*., 2019), filtered out sites containing more than 50% missing data and converted them into count files using the `cflib` package (Schrempf *et al*., 2016). The final sequences contained *∼* 400 sites. For the neutrality analyses (Tajima’s D, HKA), we also used 5-kb upstream (*∼* 400 sites) and 10-kb downstream (*∼* 900 sites) regions that are known to be neutral. The data analysis pipeline is available in the supplementary repository (https://github.com/sb2g14/PoMoBalance).

## 3 Results

### 3.1 Validation Analysis for PoMoSelect and PoMoBalance

To validate the implementations of PoMoSelect and PoMoBalance, as depicted in Figure 3, we employ the Simulation-Based Calibration procedure implemented in RevBayes (Talts *et al*., 2020). In our study, we evaluate both the PoMoSelect model with DS proposed previously by Borges *et al*. (2022b) and the model that incorporates directional and balancing selection (PoMoBalance), as outlined in Section 2.1.

In Figure 3, we conduct Simulation-Based Calibration for four PoMo functions in both reversible and non-reversible implementations, simulating the trees with five taxa and a uniform topology. The markers in the figure represent coverage probabilities for various parameters, including tree branch lengths (blue), fitnesses (*φ*, orange), nucleotide base frequencies (*π*, green), exhangeabilities (*ρ*, red), mutation rates (*μ*, rose) BS strengths (*β*, brown) and preferred frequencies (*B*, grey) and topology (purple). Different marker types distinguish values corresponding to different alleles or their combinations as per Table 2. Notably, nucleotide base frequencies exhibit a single coverage probability due to their origin from `dnDirichlet`. For fitnesses, they are relative by definition, with one of them always taking value of 1. Therefore, three coverage probabilities are observed instead of four. The 90 % confidence bounds for MCMC are shown by red dashed lines. The scores for topologies and branch lengths are best estimated for the non-reversible PoMoSelect, presumably it has fewer degrees of freedom, reducing the likelihood of encountering local minima during inference. Therefore, in this paper, we adhere to a combined approach using PoMoSelect for tree or tree topology estimation and PoMoBalance for estimating gBGC and BS.

Despite using a small virtual population size (*N* = 4) for computational efficiency, the majority of coverage probabilities lie within or very close to the confidence bounds, ensuring the validity of the implementations. The analysis of larger population sizes (*N* = 6) has shown equivalent performance.

### 3.2 Testing PoMoBalance on the data generated with Moran and SLiM simulators

In this subsection, we assess the performance of the PoMoBalance model using data simulated under various evolutionary scenarios with two different simulators. The details for the data generated with the first simulator, referred to as the Moran simulator, are depicted in Figure 2, Figure 4 (A), (B) and Figure 6 (A), (B), (C). In this analysis, we utilise RevBayes and our PoMoBalance implementation to simulate PoMo states from the non-reversible Moran model for generality, employing pre-selected parameter values akin to the scenario described in the previous subsection. However, in this case, we employ a custom phylogenetic tree depicted in Figure 4 (A), use only a few parameter sets (shown in Figure 2) and omit the calculation of coverage probabilities. Instead, we evaluate how far the inferred values deviate from the true values for a range of *σ* and *β*, as illustrated in Figure 5. Note that the accuracy of the inference decreases and confidence intervals increase with an increase in *σ* and *β*, but still the latter intersect the true values. In the case where *σ* = 0.05 and *β* = 1.5, we had to increase the number of sites from 10^5^ to 8 *×* 10^5^ for better convergence. The confidence intervals are the largest for *β* = 1, corresponding to the case where there is no balancing selection, leading to significant uncertainty in learning preferred frequencies, which affects other parameters. Additionally, we compare the SFS for *σ* = 0.1 and *β* = 2 in Figure 4 (B), calculated from the simulated data depicted by blue stars, with theoretical predictions derived using parameters inferred with PoMoBalance illustrated with red diamonds. The SFSs agree quite well despite slight inaccuracies in inferring parameters. The theoretical predictions are estimated numerically from the PoMo matrix in Equation (2), using the Markovian property d*P* (*t*)*/* d*t* = *P* (*t*)*Q*, where *P* (*t*) = exp(*tQ*). By matrix exponentiation at very long times (*t* = 10^6^), we obtain the stationary distribution for the PoMo states, which coincides with the SFS. Further details about stationary frequencies in the PoMoBalance model can be found in Supplementary Figure S1.

**Figure 5:**
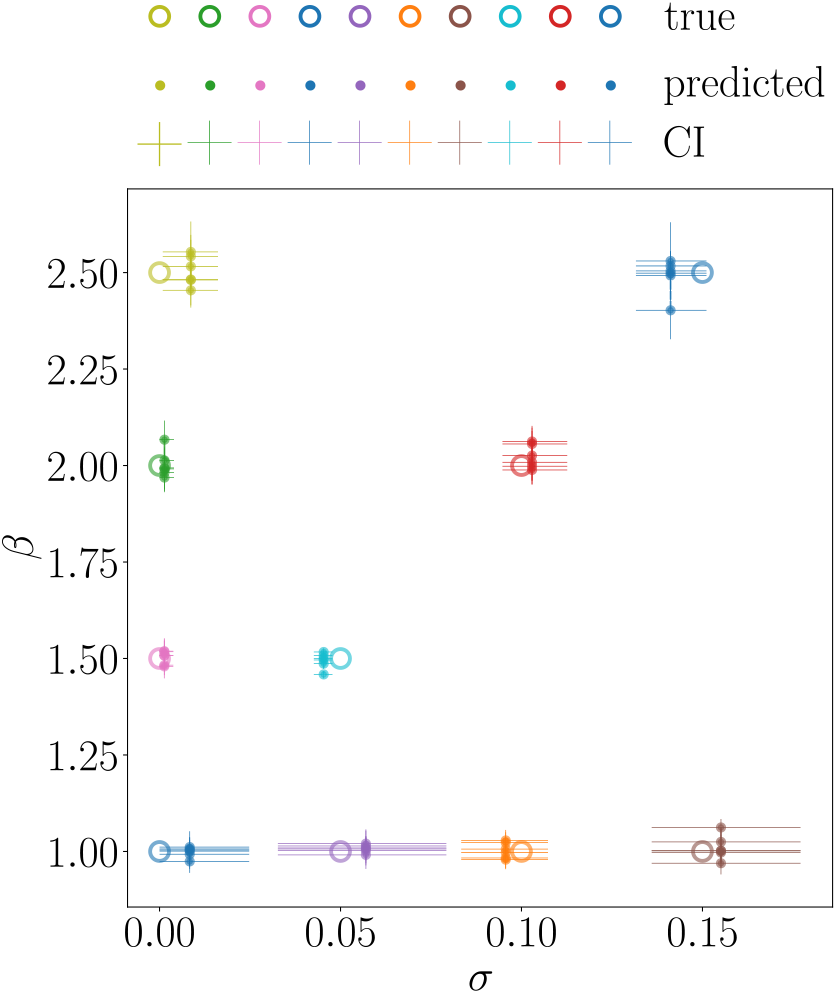
Testing PoMoBalance in a range of GC-bias rate *σ* and strength of BS *β* on the data generated with Moran model. Large open markers represent true values, smaller closed markers with error bars correspond to the mean values of posterior predictions by PoMoBalance and their 95% confidence intervals (CI) respectively.

**Figure 6:**
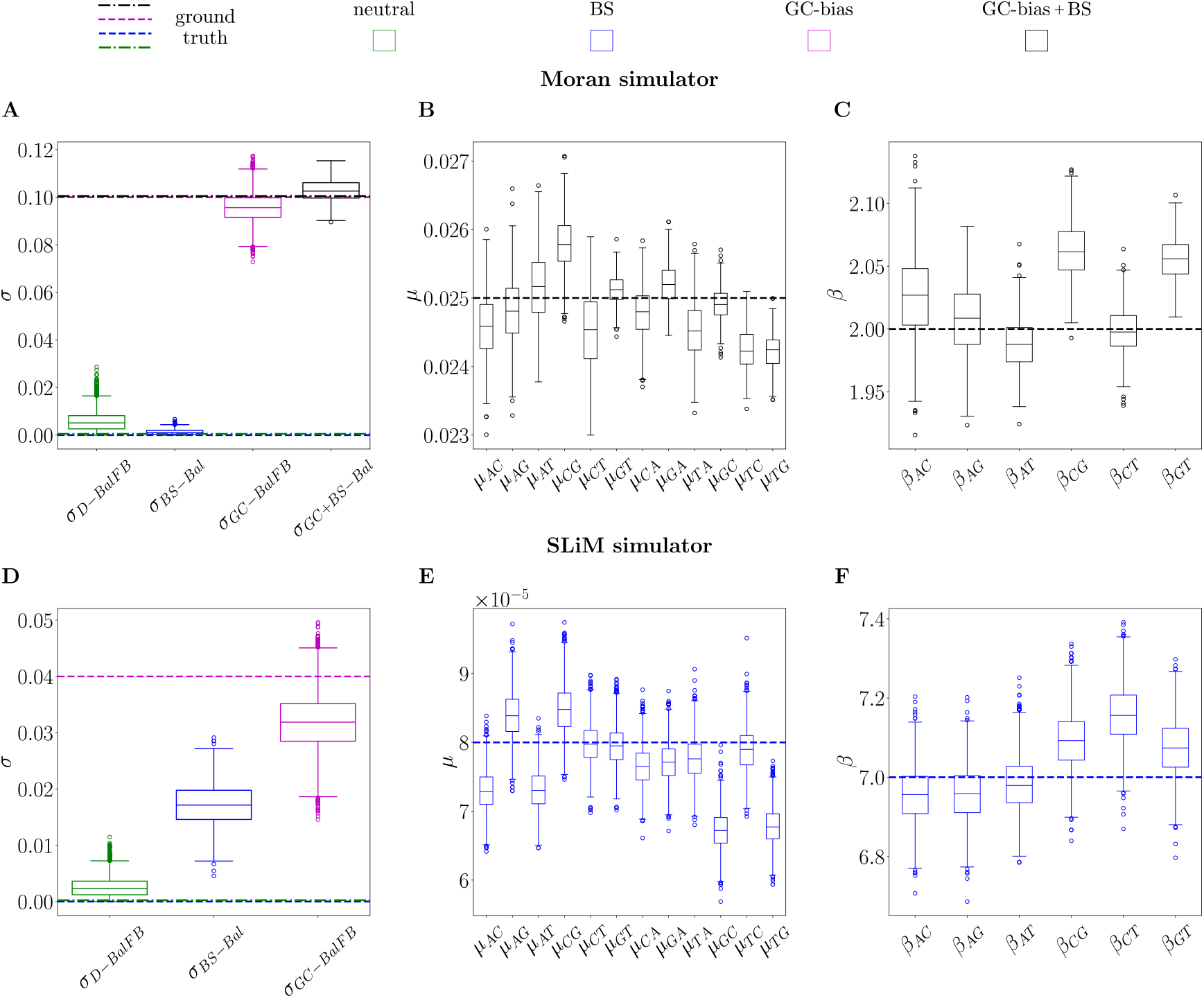
Posterior distributions of inferred parameters compared to their expected values. Subplots (A), (B), and (C) employ the Moran model simulator, in Figure 4 (A) and (B). Conversely, subplots (D), (E), and (F) use the SLiM simulator, corresponding to Figure 4 (C) and (D). Data simulations encompass four regimes: D for drift, GC for GC-biased gene conversion, BS for balancing selection, and GC+BS for the combination of gBGC and BS. Inference methods include BalFB, representing inference with PoMoBalance while fixing preferred frequencies *B*, and Bal, representing regular inference with PoMoBalance. True values are indicated by dashed and dot-dashed lines in corresponding colours. (A) Posterior plots for the GC-bias rate *σ*, with green and blue boxplots indicating simulated data in regime D inferred with BalFB and BS inferred with Bal. Magenta and black distributions correspond to regime GC inferred with BalFB and GC+BS inferred with Bal. (B) Estimates for mutation rates, and (C) strengths of BS in the simulation scenario GC+BS. (D) Posterior plots for SLiM data inference in three simulation regimes D (green), BS (blue) and GC (magenta), analogous to (A), indicating the GC-bias rate *σ*. (E) Estimates for mutation rates and (F) strengths of balancing selection corresponding to the BS simulation scenario in SLiM.

Figure 6 (A), (B) and (C) depict boxes and whiskers of the posterior distributions derived from MCMC inference with the data simulated with the Moran model. The data is simulated under four evolutionary regimes: D for neutral mutations or drift (depicted in green), GC for GC-biased gene conversion (gBGC, in magenta), BS for balancing selection (in blue), and GC+BS for the combination of gBGC and BS (in black). We plot the boxes alongside the ground truth parameters (dashed for gBGC and BS, dot-dashed for neutral and gBGC+BS) for comparison. Refer to Supplementary Table S1 for posterior means and confidence intervals for selected points.

Figure 4 (B) illustrates the SFS for the last case. In the estimation of the posterior in all cases, we discard the MCMC burn-in period.

Within the box plots in Figure 6 (A), we display estimates for the GC-bias rate in all four regimes, which align well with the true values. Mutation rates are shown in Figure 6 (B), and BS strengths are depicted in Figure 6 (C) focusing solely on the GC+BS regime for brevity. Posterior plots for preferred frequencies are not presented due to spike-like distributions as MCMC chains converge to the true values 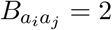 during the burn-in period. This corresponds to the BS peak in the Figure 4 (B) inset.

In Figures 4 (C), (D) and 6 (D), (E), (F), we utilise the evolutionary simulation framework SLiM proposed by Haller and Messer (2019). For this simulation, we employed the great apes tree in Supplementary Figure S2, implementing heterozygote advantage with SLiM (see Section Supplementary Material 3 for details). The tree inferred with RevBayes in Figure 4 (C) is comparable to the simulated tree, with posterior probabilities at each node equal to 1. The SFS in Figure 4 (C) is extracted from the data and features a well-distinguished peak that is effectively captured by the inference.

In SLiM simulations, we implemented three regimes (D, GC, and BS). The posterior distributions for GC-bias rate in these regimes are illustrated in Figure 6 (D). We obtain reasonable estimates in the D and GC regimes, but in the BS regime, *σ* is overestimated. This occurrence is due to the challenge of distinguishing *σ* and *π* for small virtual populations. While not easily discernible in the mutation rates presented in Figure 6 (E), it becomes apparent when examining the inferred nucleotide base frequencies *π* (refer to Supplementary Table S2). Increasing the virtual PoMo size to *N* = 20 resolves this problem partially resulting in much lower *σ*_*BS*−*Bal*_=0.008. In this analysis, our focus is on the estimation of BS strength, which shows promising results in Figure 6 (E). The preferred frequencies are also inferred accurately, similar to the Moran simulator.

Additionally, in Table 4, we present scaled scores obtained from tests conducted with MuteBaSS (HKA_trans_, NCD, NCD_opt_, NCD_sub_) and MULLET (T_1trans_, T_2trans_) (Cheng and DeGiorgio, 2019). The scores for summary statistics and likelihood-based methods were calculated using the sliding windows approach, while our method is evaluated through the logarithm of the Bayes Factor (BF). The data was generated via SLiM, similarly to Figure 4 (C), Figure 6 (D), (E) and (F) under drift, gBGC and BS regimes. For the details of the calculations please refer to Section Supplementary Material 4.

**Table 4:** Scaled by the scores calculated in the neutral case tests run with MuteBaSS (HKA_trans_, NCD, NCD_opt_, NCD_sub_) and MULLET (T_1trans_, T_2trans_) (Cheng and DeGiorgio, 2019), obtained by averaging the scores in sliding window analyses with optimal window sizes and a shift of 10 nucleotides, vs log(BF) calculated from PoMoBalance inference. The data were generated with SLiM on the tree shown in Figure 4 (C) under neutral conditions, with gBGC or BS.

| Scaled<br>Score<br>SLiM Data | $\ HKA_{trans}\ $ | $\ NCD\ $ | $\ NCD_{opt}\ $ | $\ NCD_{sub}\ $ | $\ T_{1trans}\ $ | $\ T_{2trans}\ $ | $\ \log(BF)\ $ |
| --- | --- | --- | --- | --- | --- | --- | --- |
| drift | 1.0 | 1.0 | 1.0 | 1.0 | 1.0 | 1.0 | 1.0 |
| GC-bias | 0.07 | 1.0018 | 1.003 | 1.004 | 1.01 | 0.94 | 1.01 |
| BS | 12.68 | 1.0024 | 1.012 | 1.057 | 2.26 | 4.44 | 146.05 |

The strongest evidence of BS is indicated by our method (log(BF)), followed by HKA_trans_ and T_2trans_. However, the scores of HKA_trans_ are highly dependent on the window sizes. Please note that these results must be interpreted with caution, as the scores are calculated for different approaches operating on different scales.

### 3.3 Detection of Balancing Selection in *Drosophila erecta*

In this analysis, we examine sequences derived from experimental genomic data of various *Drosophila* subspecies. We specifically explore the example of sexual dimorphism in the t_MSE_ gene region, featuring the *tan* gene observed in *Drosophila erecta* females, as studied by Yassin *et al*. (2016). Table 5 presents the results of Tajima’s D (Tajima, 1989), HKA-like (Begun *et al*., 2007) and HKA_trans_ (Cheng and DeGiorgio, 2019) tests indicating the potential presence of BS in the t_MSE_ region in contrast to neutral sequences 5-kb upstream and 10-kb downstream from the region.

**Table 5:** Results of Tajima’s D and HKA-like tests includes the number of polymorphic sites (Pol) between dark and light *Drosophila erecta* lines and divergent (Div) sites between both *erecta* lines and *Drosophila orena* in the t_MSE_ region, along with two neutral regions. The HKA_trans_ method is performed with MuteBaSS on *Drosophila erecta* (dark and light variants), *melanogaster* and *simulans* by averaging scores within 700-nucleotide windows with a step size of 10 nucleotides.

| Gene region | Tajima's D | Pol | Div | Pol/Div | HKA<sub>trans</sub> |
| --- | --- | --- | --- | --- | --- |
| t<sub>MSE</sub> | 3.99 | 51 | 28.5 | 1.78 | 0.031 |
| 5-kb upstream | -1.1 | 40 | 51.9 | 0.77 | $-6.5 \times 10^{-5}$ |
| 10-kb downstream | 0.88 | 32 | 33.5 | 0.95 | -0.175 |

The conclusion is drawn from a significant elevation of Tajimas D in the region of interest. Regarding the HKA-like test, we observe a notably higher proportion of polymorphic sites (Pol) between dark and light *Drosophila erecta* lines compared to divergent (Div) sites between both *erecta* lines and *Drosophila orena*, a closely related species to *erecta*. This increased polymorphism suggests the presence of BS. However, the chi-square test performed on these short sequences does not yield a significant result. In Yassin *et al*. (2016), the test is conducted on longer sequences containing the t_MSE_ region and leads to a significant result. The HKA_trans_ method is executed using MuteBaSS on *Drosophila erecta* (dark and light variants), *melanogaster* and *simulans*. Negative scores for the upstream and downstream regions indicate the absence of BS, unlike the positive score for the t_MSE_ region, confirming the presence of BS.

We begin the inference with PoMoSelect to determine the tree and the level of gBGC in *Drosophila* subspecies. We analyse t_MSE_ region in *Drosophila erecta* dark and light as well as *santomea, yakuba, melanogaster* and *simulans*. The tree topology obtained with PoMoSelect, as shown in Figure 7 (left), closely resembles the topology obtained by Yassin *et al*. (2016) using the multispecies coalescent method.

**Figure 7:**
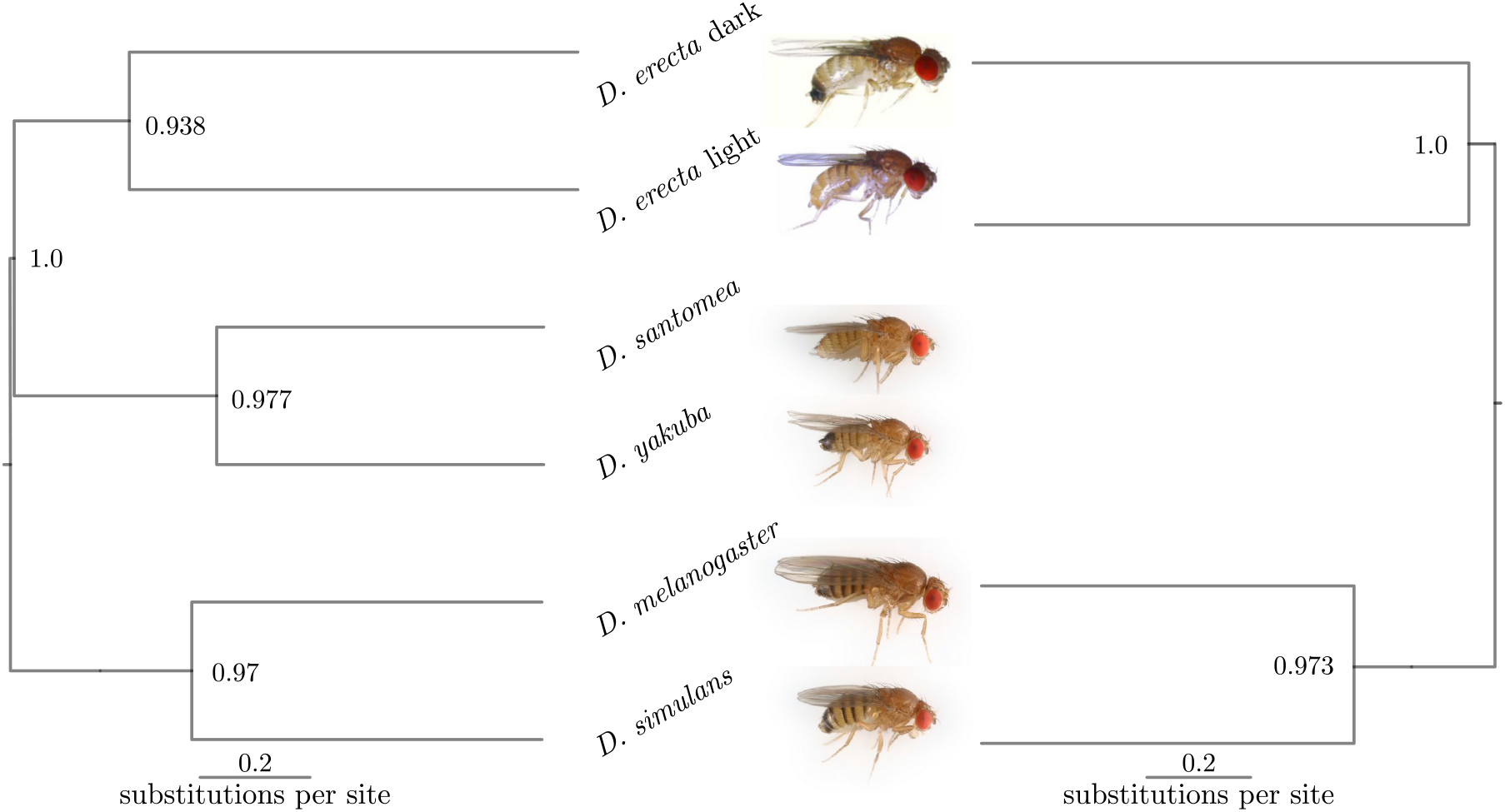
Phylogenetic tree inferred from the sequencing data obtained in the *t*_MSE_ region across six (left) and four (right) subspecies of *Drosophila*. Posterior probabilities are indicated at the nodes. Images of *D. santomea, yakuba, melanogaster* and *simulans* are credited to Darren Obbard, while those of *D. erecta* are reproduced from Yassin *et al*. (2016) under Creative Commons licence 4.0.

The gBGC rate *σ*_*Sel*_, inferred with PoMoSelect alongside the tree in Figure 7 (right), is shown in Figure 8 (A) with green box plot, and it is quite low, as observed in experiments (Robinson *et al*., 2014). Refer to Supplementary Table S3 for the inferred parameters and Effective Sample Sizes (ESS). The black box plots in Figure 8 show the posterior distributions of the parameters inferred with PoMoBalance for four *Drosophila* subspecies, namely *D. erecta* dark and light, *melanogaster* and *simulans*. Here we discard sequences of *D. santomea* and *yakuba* since they introduce noise into BS detection due to low numbers of individuals in the dataset, while still acceptable for PoMoSelect analysis. The results for all subspecies are presented in the Supplementary Material, Figures S3 and S4.

**Figure 8:**
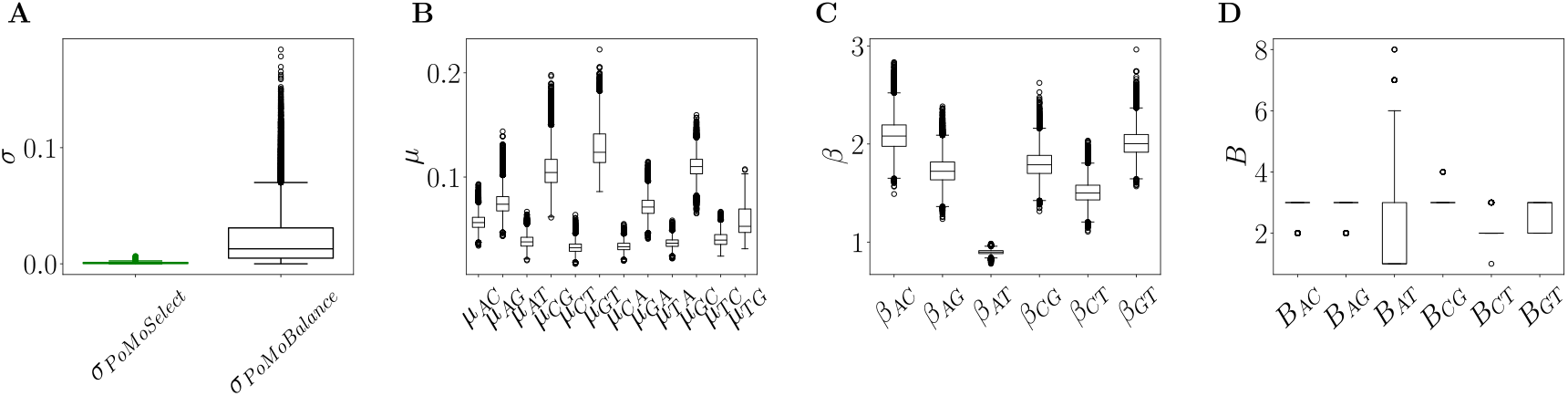
Posterior distributions derived from experimental data extracted from the *t*_MSE_ region of six subspecies, as shown in Figure 7 for PoMoSelect inference, and four *Drosophila* subspecies, namely *D. erecta* dark and light, *melanogaster* and *simulans* for PoMoBalance inference. The corresponding SFS for the PoMobalance is presented in Figure 9. (A) Estimated rates of gBGC with PoMoSelect in green and PoMoBalance in black. (B) Mutation rates, (C) strength of BS and (D) preferred frequencies for BS, all inferred using PoMoBalance.

The posterior distribution for *σ*_*P oMoBalance*_ in Figure 8 (A), inferred with PoMoBalance, is much wider than those for *σ*_*P oMoSelect*_ as it is challenging to detect GC-bias and BS simultaneously.

Thus, we advocate a mixed approach by running PoMoSelect and PoMoBalance in parallel to get more accurate estimates. For example, we learn the tree topology from PoMoSelect and then fix the estimated topology for PoMoBalance analysis. The mutation rates in Figure 8 (B) show great convergence and ESS *>* 200 for all MCMC chains. The presence of BS is detected in most of the spectra, indicated by *β >* 1 in Figure 8 (C), while for *β*_*AT*_, we observe purging of selection, indicated by *β <* 1. The preferred frequencies in Figure 8 (D) coincide or are not far away from the positions of BS peaks in the experimental SFS as shown in Figure 9.

**Figure 9:**
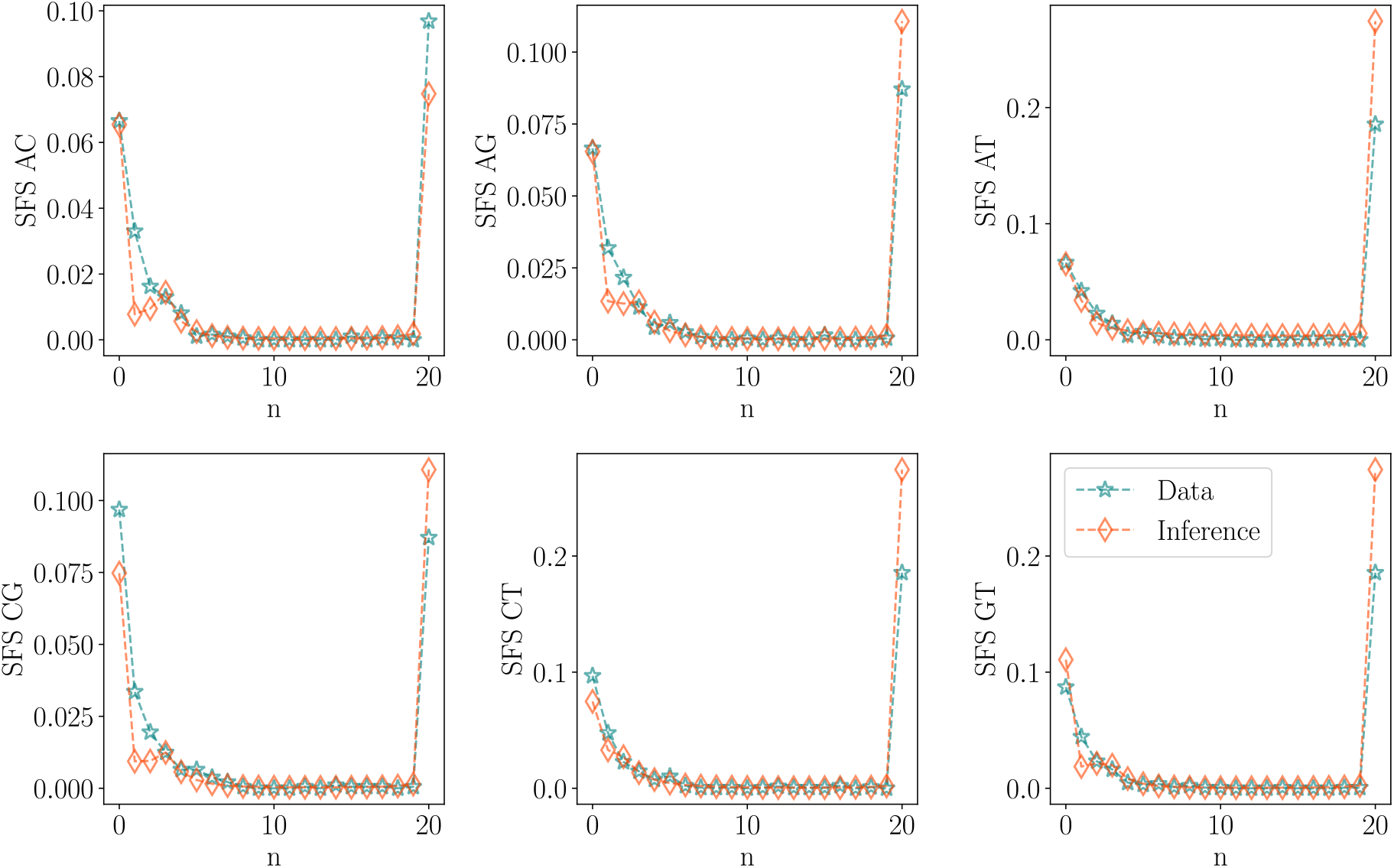
SFS representation for the t_MSE_ region corresponding to the PoMoBalance analysis in Figure 8 for four subspecies of *Drosophila*, depicted in blue stars, compared with the inferred SFS indicated by red diamonds.

We performed all analyses using the UK Crop Diversity: Bioinformatics HPC Resource and the parallel implementation of RevBayes with 24 parallel processes. The computational time was 85 hours for PoMoSelect (6 subspecies, each containing 6-25 individuals) and 118 hours (4 subspecies, each containing 6-25 individuals) for PoMoBalance to analyse the t_MSE_ region. For comparison, multispecies coalescent analysis for 2 species with introgression but without BS would take 5 days (Flouri *et al*., 2020).

## 4 Discussion

Our study validated the implementations of PoMoSelect and PoMoBalance through SBC in Subsection 3.1. Additionally, we conducted a diverse set of tests using data generated from both our custom simulator, based on the Moran model, and the evolutionary simulation framework SLiM in Subsection 2.1 (Haller and Messer, 2019). The PoMos demonstrated notable adaptability, particularly in the context of inferring data simulated via SLiM, which incorporates more complex evolutionary dynamics than the Moran model.

While SLiM, grounded in the Wright-Fisher model, shares similarities with the Moran model, it introduces additional complexities such as genetic recombination, population demography (changes in population sizes), and diploid organisms with intricate interactions between drift and heterozygote advantage. Despite these challenges, PoMoBalance performs well in locating balancing selection polymorphic peaks. To align SLiM diploids with PoMos, we treated them as two haplotypes in PoMos.

Notably, while overestimating the GC-bias rate, PoMoBalance excelled in identifying preferred frequencies, specifically in the middle of the SFS, corresponding to heterozygote advantage in SLiM. This represents a unique advantage compared to previous methods, which, while suggestive of the presence of balancing selection, cannot pinpoint specific combinations of alleles, strengths, and preferred frequencies of balancing selection. It is important to acknowledge potential correlations between *β* and *σ*, which limits their inference. To address this, we advocate for incorporating extra moves into the MCMC, as discussed in Subsection 2.2. The comparative analysis with MuteBaSS and MULLET indicates that our method demonstrates the strongest evidence of BS for data involving the heterozygote advantage. However, this result must be interpreted with caution since we assess the performance of our method using the BF approach, while we derive averaged statistics for the other methods (see Section Supplementary Material 4).

In Subsection 3.3, we applied PoMoSelect and PoMoBalance to analyse experimental genomic data from *Drosophila erecta*, specifically focusing on the t_MSE_ region known to exhibit balancing selection (Yassin *et al*., 2016). Our application of PoMos reproduced previous insights by Yassin *et al*. (2016) into the phylogenetic relationships among *Drosophila* subspecies.

Note, that the outcomes of the inference for CG-bias rate and mutation rates are presented in terms of the virtual PoMos population sizes, which typically differ from the actual population sizes. To accurately reflect the actual population dynamics in *Drosophila*, it is necessary to map the values of *μ, σ, β* and *B* from virtual PoMos size to effective population size (see Supplementary Material 2). This mapping results in substantially reduced parameter values for *σ* and *μ*, as found by Borges *et al*. (2019), given the large effective population sizes characteristic of *Drosophila* (Kelley *et al*., 2005). The mapping for the preferred frequency is relatively straightforward, and we plan to propose a mapping for the BS strengths and the non-reversible coefficients in future research.

Through PoMoBalance analysis, we detect BS in the majority of allele combinations, in contrast to the absence of BS peaks in neutral regions. Additionally, we observe the purging of selection for AT alleles, signifying the removal of polymorphisms at a rate higher than expected under neutral conditions. While this discovery showcases the flexibility of our method, interpreting its biological implications is challenging. Moreover, such interpretation might be unnecessary, as the mean value for *β*_*AT*_ is only slightly smaller than 1, indicating neutrality expectations and suggesting a relatively weak effect.

## 5 Conclusion

We incorporated the PoMoBalance model, a generalised form of PoMos capable of detecting BS, into RevBayes, a widely used phylogenetic software based on Bayesian inference. This integration enriches the resources available to researchers engaged in phylogenetic analysis, providing a robust framework for precise species tree inference and concurrent parameter estimation. Notably, our implementation allows for the estimation of balancing selection, including preferred frequencies and specific alleles under selection, while also disentangling it from other forms of selection. PoMoBalance exhibits versatility in capturing various selection types, including purging selection, observed when the level of observed polymorphisms is lower than expected via genetic drift and directional selection. These effects may arise from a combination of dominance effects, such as underdominance, or purifying selection in the context of background selection, etc.

In general, we provide a comprehensive framework to use PoMos for the estimation of phylogenetic trees, GC-bias and BS. The approach involves several key steps. First, we employ the PoMoSelect to estimate tree topology, GC-bias rate, and mutations. Subsequently, we use PoMoBalance to estimate all parameters, allowing branch lengths to vary while maintaining a fixed topology learned from PoMoSelect. It is worthwhile to validate the results by comparing the inferred values with PoMoBalance estimates that include a fixed GC-bias rate learned from PoMoSelect. The selection of the best candidates is based on the agreement between the inferred SFS and that estimated from the data. Lastly, in this framework, PoMoBalance is selectively applied to regions that are likely under balancing selection, such as the MHC locus in *Homo sapiens*.

The adaptability and versatility of PoMos address a need in the analysis of complex genomic datasets since our framework provides accurate phylogenetic inferences across multiple timescales and demonstrate potential for application in genome-wide scans through the parallel inference of multiple genomic regions. The other benefit of PoMos is scalability in terms of the number of species; it is capable of handling dozens of species (Borges *et al*., 2022a). In future, we aim to investigate additional genomic factors intertwined with balancing selection, with a specific focus on exploring the role of linkage disequilibrium and its impact on the detection of BS.

## 6 Software Availability

The software RevBayes (Höhna *et al*., 2016; Hohna *et al*., 2017; Höhna *et al*., 2018) is available at https://revbayes.github.io/. PoMoBalance tutorial at https://revbayes.github.io/tutorials/pomobalance/.

## 7 Data Availability

The data and the code for PoMoBalance analysis concerning Simulation-Based Calibration, Moran simulator and SLiM are available via GitHub (https://github.com/sb2g14/PoMoBalance). The sequencing data for *Drosophila erecta* and *orena* used in the analysis was previously published by Yassin *et al*. (2016), the data for multiple individuals of other related subspecies of *Drosophila* was obtained via NCBI BLAST (https://blast.ncbi.nlm.nih.gov/Blast.cgi).

## 8 Acknowledgments

We thank Sebastian Höhna, Amir Yassin, Valeria Montano, Dominik Schrempf and Ben Haller for helpful discussions. This work was supported by Biotechnology and Biological Sciences Research Council (BBSRC) [BBW00768/1 and BB/Y513842/1]. The authors acknowledge Research Computing at the James Hutton Institute for providing computational resources and technical support for the “UK’s Crop Diversity Bioinformatics HPC” (BBSRC grants BB/S019669/1 and BB/X019683/1), use of which has contributed to the results reported within this paper.

## Supplementary Material 1 Stationary Distribution and Reversibility in PoMoBalance

The stationary distribution *ψ*provides the opportunity to investigate the long-term behavior of the interplay between mutational bias, genetic drift, directional, and balancing selection on population diversity. In the biallelic case, the Moran dynamic exemplifies a birth-and-death process, known for its reversibility. Consequently, we obtained the stationary distribution by initially formulating the detailed balance equations. To simplify the notation, we redefine the state *{na*_*i*_, (*N* − *n*)*a*_*j*_*}* to only represent the frequency of the *a*_*i*_ allele: i.e., *{n}*.

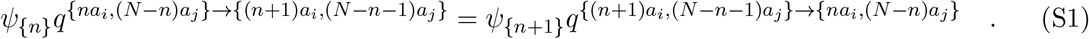

The detailed balance equations allow us to derive the following recursive formula, which is employed to obtain the stationary quantities for both fixed and polymorphic states

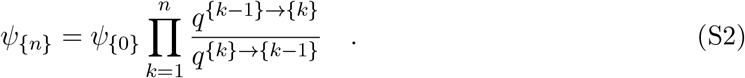

If we set *n* = *N*, the recursive formula becomes

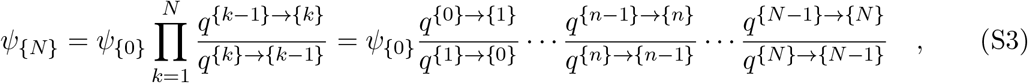

from which by considering the rates of the process defined in the rate matrix *Q* in Equation (2), we find the normalized stationary quantities for the fixed states

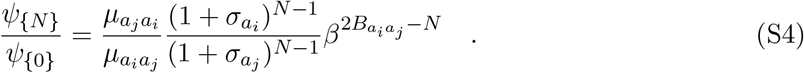

An interesting aspect is that the differentiated impact of BS in the fixed states disappears when the balanced frequency sits in the middle of the frequency spectrum (i.e., 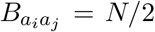). By applying the Kolmogorov criterion to each closed chain in the PoMoBalance model described with Equation (2) we ensure that reversibility is satisfied when 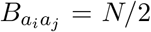 and breaks in all other cases.

The stationary distributions *ψ*_*{n}*_ may be multiplied by any arbitrary constant without affecting the final result thanks to the normalisation condition. Thus, we are safe to assume that we could set

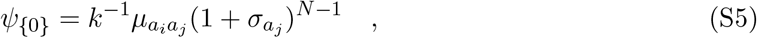

where *k* is obtained from the normalisation condition 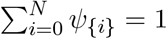. Then from equation (S4) we find

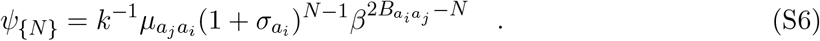

Similarly to the fixed states, the stationary measures for the polymorphic states can be derived using the recursive formula in equation (S2)

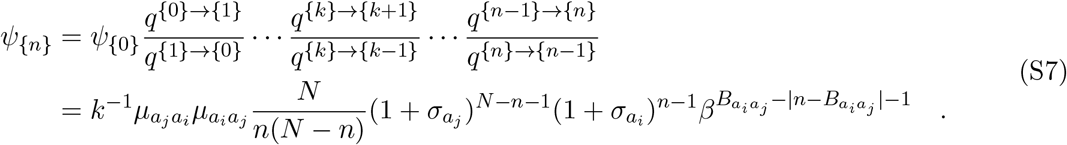

This solution clearly illustrates the contribution of mutational bias, genetic drift, directional selection, and BS to the frequency of polymorphic states. As expected, the BS term is highest when 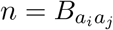 and decays in the direction of the boundary states. This feature becomes evident when we compare the stationary distribution with and without the effect of BS in Figure S1.

**Figure S1:**
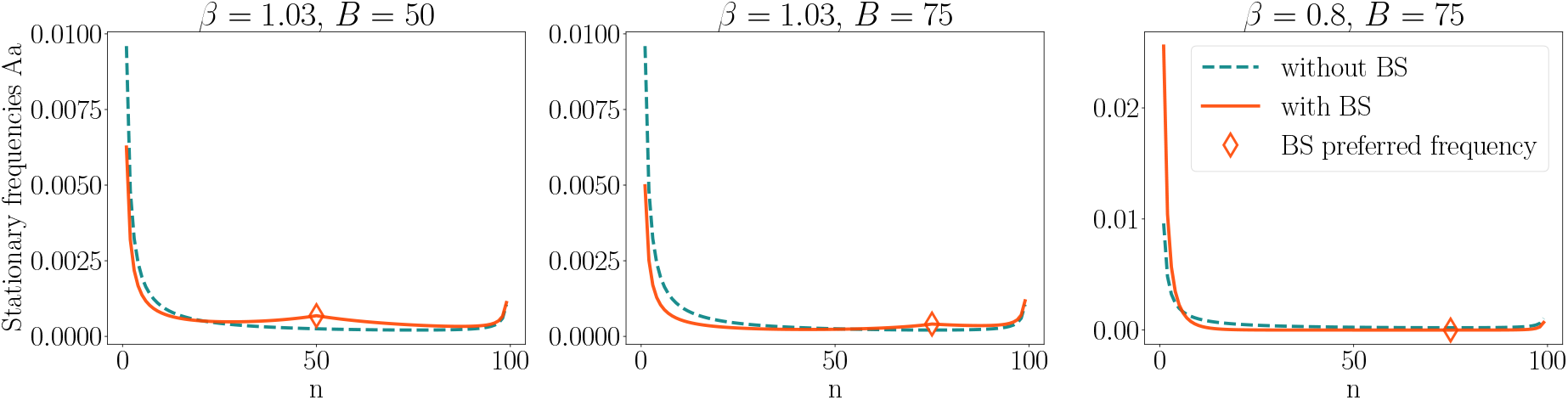
The plots depict the stationary distribution of a population of *N* = 100 individuals, and a biallelic locus with alleles *A* and *a* that evolves under under mutational bias (*μ*_*Aa*_ = 0.02 *> μ*_*aA*_ = 0.01), directional selection (*σ*_*A*_ = 0.01 *> σ*_*a*_ = 0.0) and three regimes of BS. Here we present frequencies in the range [1, *N* − 1] to avoid very high tails that dominate the BS peak.

Because we are interested in modelling BS, we have been assuming that 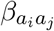 acts to maintain diversity at a certain frequency 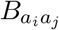. Mathematically speaking, we have been assuming that 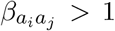. However, an interesting behaviour emerges when 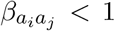. In this case, the BS term acts to purge variation more than what is already expected by genetic drift and directional selection, as shown in Figure S1. We refer to this regime as purging selection.

We normalize the obtained stationary quantities obtained in equations (S5), (S6) and (S7) to sum up to 1. The stationary distribution normalization constant is

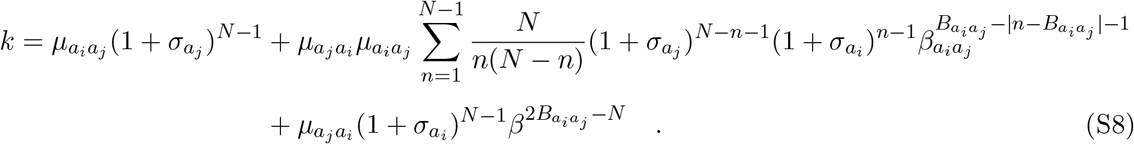

## Supplementary Material 2 Virtual population size in PoMoBalance

In PoMos, we operate with virtual population sizes for computational efficiency. Although the effective population size does not directly reflect the exact mutation rates, gBGC coefficients, and balancing selection coefficients, it provides a means to assess the relative strength of these effects. Mapping these coefficients to actual population sizes is feasible but depends on the specific PoMos utilised in the analysis.

For instance, Borges *et al*. (2019) detailed in Appendix A the mapping between the virtual sizes of reversible PoMoSelect for two populations, *A* and *A*^*t*^ with respective sizes *N* and *M*. In the case of reversible PoMoBalance, the scaling would be similar, utilising SFS in Equations (S5), (S6) and (S7) where *σ* and *β* would need to be scaled together. The preferred frequency scales as 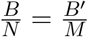, where *B* represents the frequency of population *A*, and *B*^*’*^ is the frequency of population *A*^*’*^.

Regarding non-reversible PoMos, the mapping of the preferred frequency remains unchanged, while the other coefficients require numerical mapping of solutions *P* (*t*) = exp(*tQ*) for population *A* and *P* ^*‘*^(*t*) = exp(*tQ*^*’*^) for *A*^*’*^. This aspect requires further investigation in future research endeavors.

### Supplementary Material 3 Simulations with SLiM

The original scripts for SLiM simulations can be found in the supplementary repository (https://github.com/sb2g14/PoMoBalance). We ran nucleotide models, simulating 10^5^ genomic sites with drift only using ‘`initializeMutationTypeNuc(“m1”, 0.5, “f”, 0.0)`’, drift+gcbias using ‘initializeGeneConversion(0.3, 1500, 0.80, 1.0)’, drift+heterozygote advantage with ‘`initializeMutationTypeNuc(“m2”, 1.1, “f”, 0.1)`’, where the coefficient 1.1 simulates overdominance. We set the mutation rate to 10^−6^ and the recombination rate to 10^−5^. These rates are higher than physiological ones for computational purposes, but they work well for the purposes of our analysis.

**Figure S2:**
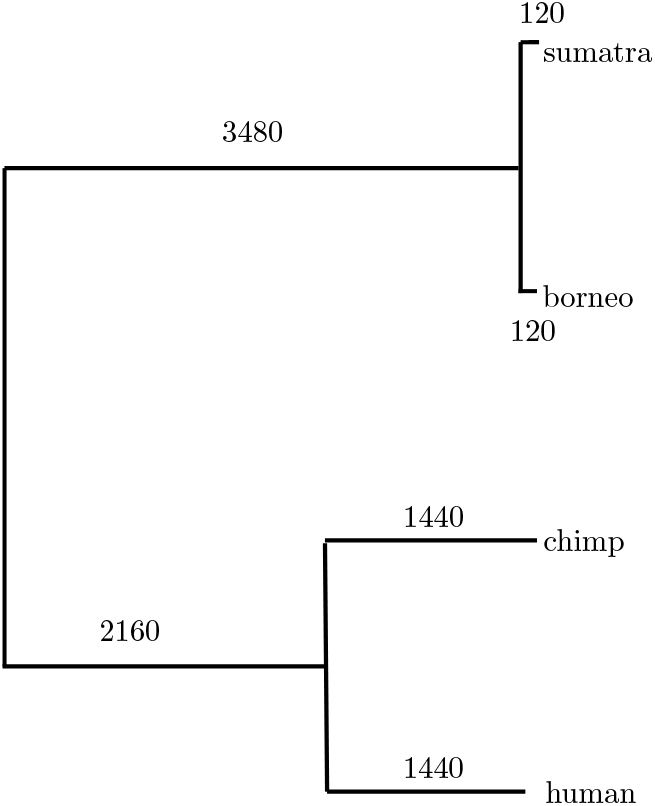
Phylogenetic tree simulated with SLiM, the inferred tree is presented in Figure 4 (C), here the branch lengths are expressed in the simulated generations.

We initialised a population of 2000 individuals of *Homininae* and evolved them for 10000 populations as a burn-in step. Then we split them into 1000 of *Hominini* and 1000 of *Gorillini*. Following the numbers of generations shown in Figure 1 we end up with *Orangutan sumatra, Orangutan borneo, chimp* and *human*, each containing 500 individuals.

Finally, the ancestral sequences (`.FASTA`) and polymorphic data (`.VCF`) are written out as output for each population.

Note that in the inference with the data simulated with SLiM, as shown in Table S2, the BS strengths *β* for the neutral case and GC-biased case are significantly underestimated (the most are 0.7 instead of 1). This is presumably due to noise in the simulations, especially when SLiM diploids are mapped to small population sizes in PoMos. Interestingly, with an increase in the population size, such misspecification is reduced, and for *N* = 20, we obtain *β* = 0.9.

## Supplementary Material 4 Model comparison

In Table 4 we strive to compare several approaches for detecting BS. Here, we utilise data generated with SLiM to produce Figure 6 For MuteBaSS (HKA_trans_, NCD, NCD_opt_, NCD_sub_) we determine the optimal window size from the sliding window analysis, which yields corresponding scores for each window. We consider a range of window sizes (refer to Table S4 for drift, Table S5 for gBGC and Table S6 for BS) with a small step size (10 bases) to ensure a sufficient number of scores 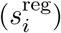 in each run. in each run. The resulting scores, presented in Tables S4, S5 and S6, represent averages of all 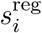 within each run:

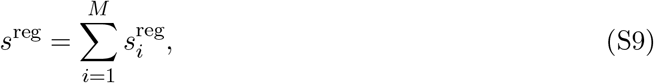

where *M* is the total number of calculated scores in each sliding window run, and reg denotes the regime under which the data was obtained (drift, gBGC, or BS). The scaled scores provided in Table 4 are calculated by scaling each averaged score by the neutral case 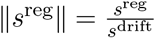. The optimal window size is selected from all scores in Table S5 where ∥*s*^gBGC^∥ is closest to 1 (except for HKA_trans_, which is the only one testing for selection), indicating no BS in the data generated with gBGC, while ∥*s*^BS^∥ simultaneously reaches its maximum.

The scaled scores for MULLET (T_1trans_ and T_2trans_) (Cheng and DeGiorgio, 2019) are calculated similarly to MuteBaSS, with the only difference being the omission of the step size. In the T statistics, the region assessed at each step centers on a test-informative site and includes a set number of informative sites both upstream and downstream, rather than employing a fixed physical window size based on the number of nucleotides. It’s worth noting that for a large number of informative sites (over 500 or, in some cases, 1000), the T statistics always returns NAN.

The log Bayes Factors in PoMos analysis are calculated from the difference of marginal log-likelihoods in each regime

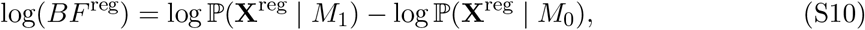

where *M*_0_ represents the PoMoSelect model, while *M*_1_ represents PoMoBalance. **X**^reg^ is the data generated in certain regime (drif, gBGC and BS). log ℙ(**X**^reg^ | *M*_0_) and log ℙ(**X**^reg^ | *M*_1_) are corresponding log-likelihoods. In general, log(*BF* ^reg^) explains how much PoMoBalance explains data better comparing to PoMoSelect. The values for log-likelihoods in our analysis are log ℙ(**X**^drift^ | *M*_0_) = −290451.8918, log ℙ(**X**^gBGC^ | *M*_0_) = −282710.9797, log ℙ(**X** ^BS^ | *M*_0_) = −308840.628, log ℙ(**X**^drift^ | *M*_1_) = −289601.7246, log ℙ(**X**^gBGC^ | *M*_1_) = −281708.4163 and log ℙ(**X** ^BS^ | *M*_1_) = −245036.4488. Similarly to the previous cases the scaled score is 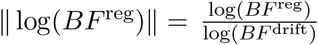.

## Supplementary Material 5 Supplementary Figures

**Figure S3:**
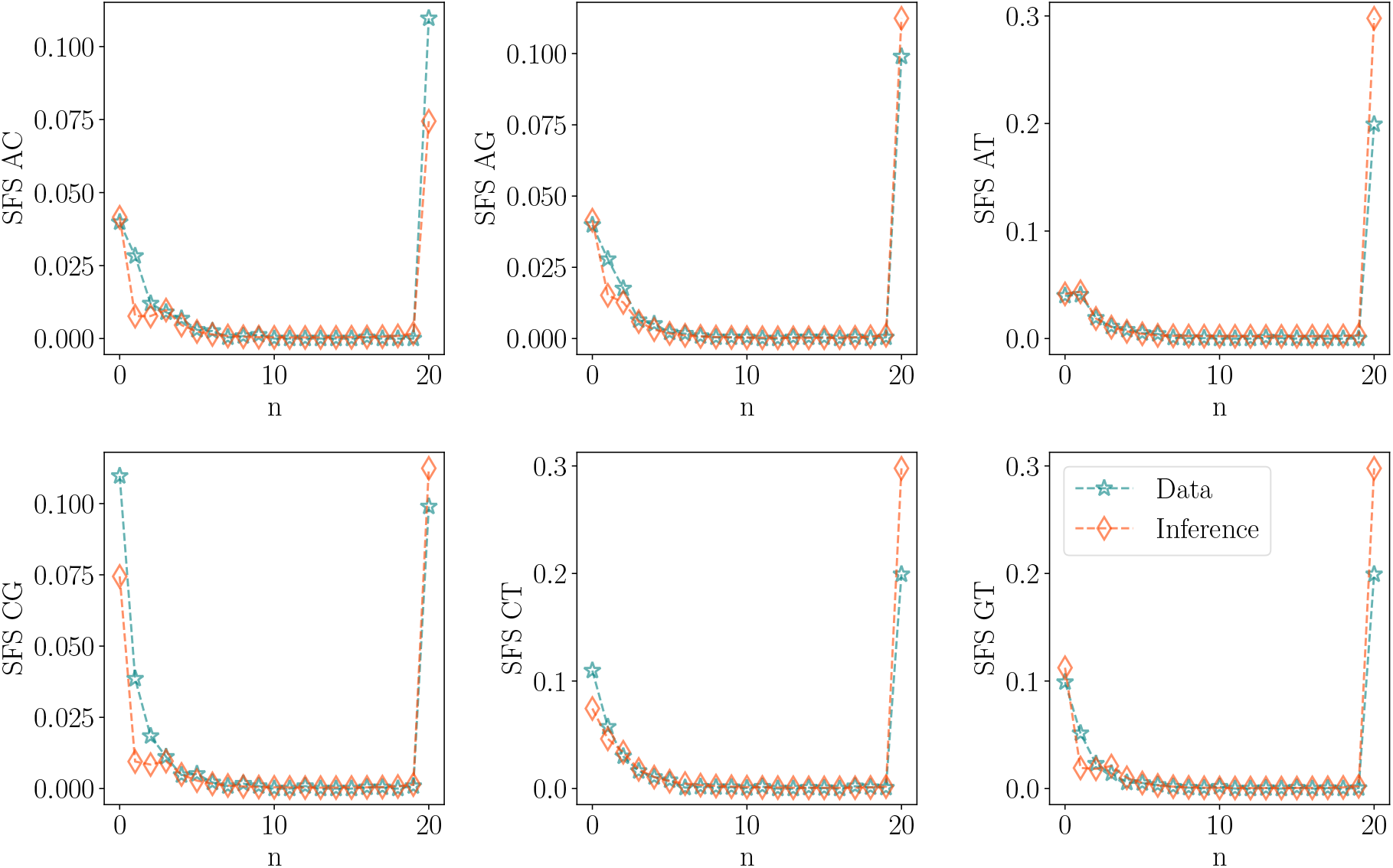
SFS representation for the *t*_MSE_ region in six subspecies of *Drosophila*, denoted by blue stars, is compared with the SFS inferred using PoMoBalance, represented by red diamonds.

**Figure S4:**
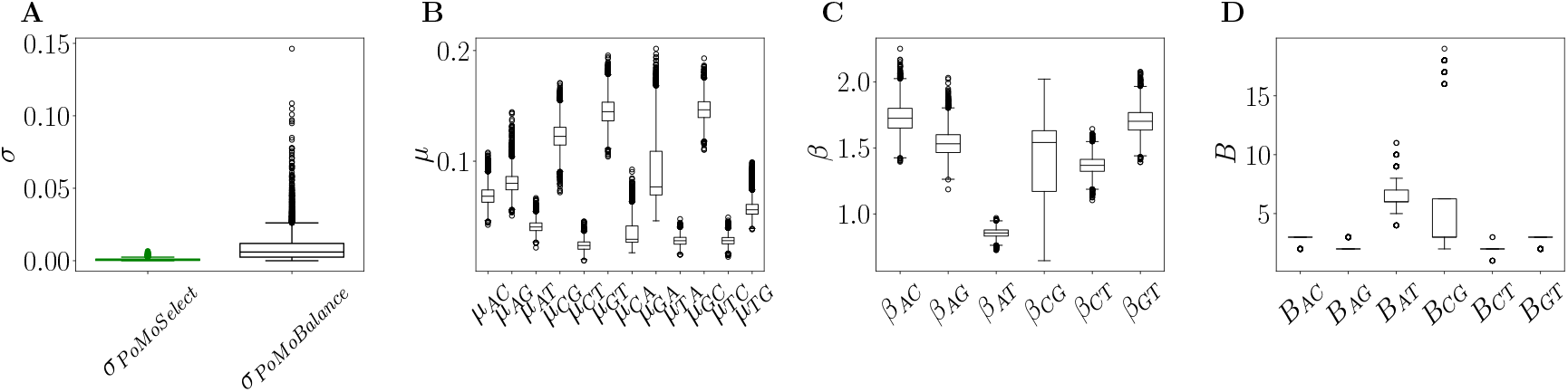
Posterior distributions derived from experimental data extracted from the *t*_MSE_ region of six *Drosophila* subspecies. The corresponding tree and SFS are presented in Figure 7 and S3. (A) Estimated rates of gBGC with PoMoSelect in green and PoMoBalance in black. (B) Mutation rates, (C) strength of BS and (D) preferred frequencies for BS, all inferred using PoMoBalance.

## Supplementary Material 6 Supplementary Tables

**Table S1:** Results of converged MCMC chain for Figure 4 (A), (B) and 6 (A), (B), (C).

|  | Drift |  |  | GC-bias |  |  |
| --- | --- | --- | --- | --- | --- | --- |
| Variable | True values | Posterior mean | 95 % credible interval | True values | Posterior mean | 95 % credible interval |
| sigma | 0 | 0.042 | [0, 0.205] | 0.1 | 0.098 | [0.0013, 0.197] |
| pi_A | 0.25 | 0.257 | [0.253, 0.261] | 0.25 | 0.249 | [0.244, 0.255] |
| pi_C | 0.25 | 0.252 | [0.247, 0.256] | 0.25 | 0.254 | [0.248, 0.259] |
| pi_G | 0.25 | 0.242 | [0.238, 0.247] | 0.25 | 0.249 | [0.244, 0.254] |
| pi_T | 0.25 | 0.249 | [0.244, 0.254] | 0.25 | 0.248 | [0.243, 0.253] |
| rho_AC | 0.1 | 0.098 | [0.095, 0.1] | 0.1 | 0.099 | [0.096, 0.101] |
| rho_AG | 0.1 | 0.1 | [0.098, 0.103] | 0.1 | 0.1 | [0.097, 0.103] |
| rho_AT | 0.1 | 0.098 | [0.096, 0.101] | 0.1 | 0.1 | [0.097, 0.103] |
| rho_CG | 0.1 | 0.102 | [0.099, 0.105] | 0.1 | 0.1 | [0.097, 0.103] |
| rho_CT | 0.1 | 0.099 | [0.097, 0.102] | 0.1 | 0.098 | [0.095, 0.1] |
| rho_GT | 0.1 | 0.101 | [0.098, 0.103] | 0.1 | 0.102 | [0.1, 0.105] |
| beta_AC | 1 | 1.039 | [0.968, 1.198] | 1 | 1.014 | [0.911, 1.098] |
| beta_AG | 1 | 1.01 | [0.876, 1.198] | 1 | 1.018 | [0.926, 1.113] |
| beta_AT | 1 | 1.006 | [0.976, 1.035] | 1 | 0.998 | [0.959, 1.042] |
| beta_CG | 1 | 0.978 | [0.945, 1.03] | 1 | 0.997 | [0.969, 1.025] |
| beta_CT | 1 | 1.003 | [0.807, 1.147] | 1 | 0.999 | [0.892, 1.077] |
| beta_GT | 1 | 1.029 | [0.959, 1.232] | 1 | 0.991 | [0.894, 1.072] |
|  | BS |  |  | GC-bias + BS |  |  |
| sigma | 0 | 0.0014 | [0, 0.004] | 0.1 | 0.0014 | [0, 0.004] |
| pi_A | 0.25 | 0.248 | [0.242, 0.253] | 0.25 | 0.254 | [0.247, 0.26] |
| pi_C | 0.25 | 0.254 | [0.249, 0.258] | 0.25 | 0.252 | [0.247, 0.258] |
| pi_G | 0.25 | 0.245 | [0.242, 0.249] | 0.25 | 0.247 | [0.243, 0.252] |
| pi_T | 0.25 | 0.253 | [0.25, 0.257] | 0.25 | 0.248 | [0.243, 0.252] |
| rho_AC | 0.1 | 0.102 | [0.099, 0.106] | 0.1 | 0.098 | [0.094, 0.101] |
| rho_AG | 0.1 | 0.101 | [0.099, 0.105] | 0.1 | 0.102 | [0.099, 0.105] |
| rho_AT | 0.1 | 0.101 | [0.098, 0.104] | 0.1 | 0.099 | [0.095, 0.102] |
| rho_CG | 0.1 | 0.1 | [0.097, 0.102] | 0.1 | 0.1 | [0.097, 0.103] |
| rho_CT | 0.1 | 0.098 | [0.095, 0.1] | 0.1 | 0.099 | [0.096, 0.101] |
| rho_GT | 0.1 | 0.101 | [0.099, 0.103] | 0.1 | 0.098 | [0.096, 0.1] |
| B_AC | 2 | 2 | [2.0, 2.0] | 2 | 2 | [2.0, 2.0] |
| B_AG | 2 | 2 | [2.0, 2.0] | 2 | 2 | [2.0, 2.0] |
| B_AT | 2 | 2 | [2.0, 2.0] | 2 | 2 | [2.0, 2.0] |
| B_CG | 2 | 2 | [2.0, 2.0] | 2 | 2 | [2.0, 2.0] |
| B_CT | 2 | 2 | [2.0, 2.0] | 2 | 2 | [2.0, 2.0] |
| B_GT | 2 | 2 | [2.0, 2.0] | 2 | 2 | [2.0, 2.0] |
| beta_AC | 2 | 1.991 | [1.934, 2.047] | 2 | 1.991 | [1.934, 2.047] |
| beta_AG | 2 | 1.982 | [1.931, 2.029] | 2 | 1.982 | [1.931, 2.029] |
| beta_AT | 2 | 1.994 | [1.958, 2.029] | 2 | 1.994 | [1.958, 2.029] |
| beta_CG | 2 | 2.067 | [2.022, 2.117] | 2 | 2.067 | [2.022, 2.117] |
| beta_CT | 2 | 2.013 | [1.98, 2.043] | 2 | 2.013 | [1.98, 2.043] |
| beta_GT | 2 | 1.969 | [1.939, 1.996] | 2 | 1.969 | [1.939, 1.996] |

**Table S2:** Results of converged MCMC chain for Figure 4 (C), (D) and 6 (D), (E), (F).

|  | **Drift** |  |  | **GC-bias** |  |  |
| --- | --- | --- | --- | --- | --- | --- |
| Variable | True values | Posterior mean | 95 % credible interval | True values | Posterior mean | 95 % credible interval |
| sigma | 0 | 0.0026 | [0, 0.0058] | 0.04 | 0.032 | [0.023, 0.042] |
| pi_A | 0.25 | 0.252 | [0.248, 0.257] | 0.25 | 0.166 | [0.16, 0.172] |
| pi_C | 0.25 | 0.249 | [0.245, 0.253] | 0.25 | 0.23 | [0.224, 0.238] |
| pi_G | 0.25 | 0.248 | [0.244, 0.252] | 0.25 | 0.302 | [0.294, 0.311] |
| pi_T | 0.25 | 0.251 | [0.246, 0.255] | 0.25 | 0.301 | [0.291, 0.311] |
| rho_AC | 0.04 | 0.041 | [0.04, 0.043] | 0.03 | 0.047 | [0.044, 0.049] |
| rho_AG | 0.04 | 0.042 | [0.041, 0.044] | 0.03 | 0.037 | [0.035, 0.038] |
| rho_AT | 0.04 | 0.041 | [0.039, 0.043] | 0.03 | 0.031 | [0.029, 0.032] |
| rho_CG | 0.04 | 0.042 | [0.04, 0.044] | 0.03 | 0.031 | [0.03, 0.033] |
| rho_CT | 0.04 | 0.039 | [0.038, 0.041] | 0.03 | 0.029 | [0.028, 0.031] |
| rho_GT | 0.04 | 0.039 | [0.037, 0.04] | 0.03 | 0.03 | [0.028, 0.031] |
| beta_AC | 1 | 0.71 | [0.692, 0.727] | 1 | 0.749 | [0.73, 0.769] |
| beta_AG | 1 | 0.719 | [0.7, 0.736] | 1 | 0.822 | [0.799, 0.845] |
| beta_AT | 1 | 0.714 | [0.696, 0.732] | 1 | 1.094 | [1.069, 1.12] |
| beta_CG | 1 | 0.7 | [0.682, 0.717] | 1 | 0.794 | [0.775, 0.814] |
| beta_CT | 1 | 0.711 | [0.693, 0.73] | 1 | 0.849 | [0.815, 0.883] |
| beta_GT | 1 | 0.733 | [0.714, 0.751] | 1 | 0.694 | [0.668, 0.718] |
|  | **BS** |  |  |  |  |  |
| sigma | 0 | 0.017 | [0.01, 0.025] |  |  |  |
| pi_A | 0.25 | 0.268 | [0.259, 0.276] |  |  |  |
| pi_C | 0.25 | 0.233 | [0.224, 0.241] |  |  |  |
| pi_G | 0.25 | 0.231 | [0.222, 0.238] |  |  |  |
| pi_T | 0.25 | 0.269 | [0.261, 0.277] |  |  |  |
| rho_AC | 0.00032 | 0.00031 | [0.00029, 0.00034] |  |  |  |
| rho_AG | 0.00032 | 0.00032 | [0.0003, 0.00034] |  |  |  |
| rho_AT | 0.00032 | 0.0003 | [0.00027, 0.00032] |  |  |  |
| rho_CG | 0.00032 | 0.00033 | [0.0003, 0.00036] |  |  |  |
| rho_CT | 0.00032 | 0.00029 | [0.00027, 0.00031] |  |  |  |
| rho_GT | 0.00032 | 0.00029 | [0.00027, 0.00032] |  |  |  |
| B_AC | 5.0 | 5.0 | [5.0, 5.0] |  |  |  |
| B_AG | 5.0 | 5.0 | [5.0, 5.0] |  |  |  |
| B_AT | 5.0 | 5.0 | [5.0, 5.0] |  |  |  |
| B_CG | 5.0 | 5.0 | [5.0, 5.0] |  |  |  |
| B_CT | 5.0 | 5.0 | [5.0, 5.0] |  |  |  |
| B_GT | 5.0 | 5.0 | [5.0, 5.0] |  |  |  |
| beta_AC | 7.0 | 6.956 | [6.814, 7.085] |  |  |  |
| beta_AG | 7.0 | 6.958 | [6.827, 7.095] |  |  |  |
| beta_AT | 7.0 | 6.981 | [6.839, 7.107] |  |  |  |
| beta_CG | 7.0 | 7.093 | [6.956, 7.228] |  |  |  |
| beta_CT | 7.0 | 7.158 | [7.027, 7.304] |  |  |  |
| beta_GT | 7.0 | 7.074 | [6.934, 7.217] |  |  |  |

**Table S3:** Results of the inference combined from 4 MCMC chains for Figure 8 and 9.

|  | **PoMoSelect** |  |  | **PoMoBalance** |  |  |
| --- | --- | --- | --- | --- | --- | --- |
| Variable | Posterior<br>mean | 95 %<br>credible interval | ESS | Posterior<br>mean | 95 %<br>credible interval | ESS |
| sigma | 0.00081 | [0, 0.0023] | 7660 | 0.022 | [0, 0.074] | 472 |
| pi_A | 0.176 | [0.139, 0.216] | 1773 | 0.422 | [0.358, 0.488] | 341 |
| pi_C | 0.292 | [0.254, 0.33] | 3696 | 0.32 | [0.278, 0.365] | 1645 |
| pi_G | 0.225 | [0.191, 0.258] | 3760 | 0.151 | [0.115, 0.193] | 241 |
| pi_T | 0.306 | [0.272, 0.343] | 4787 | 0.107 | [0.082, 0.134] | 641 |
| rho_AC | 0.028 | [0.021, 0.036] | 2950 | 0.178 | [0.135, 0.226] | 759 |
| rho_AG | 0.032 | [0.024, 0.041] | 3008 | 0.255 | [0.193, 0.331] | 2654 |
| rho_AT | 0.025 | [0.019, 0.031] | 2267 | 0.304 | [0.238, 0.379] | 1920 |
| rho_CG | 0.022 | [0.018, 0.026] | 4943 | 0.224 | [0.173, 0.28] | 1286 |
| rho_CT | 0.02 | [0.017, 0.023] | 4726 | 0.344 | [0.284, 0.409] | 2377 |
| rho_GT | 0.018 | [0.015, 0.021] | 4042 | 0.379 | [0.286, 0.485] | 406 |
| B_AC | - | - | - | 2.995 | [3.0, 3.0] | 2375 |
| B_AG | - | - | - | 2.86 | [2.0, 3.0] | 1315 |
| B_AT | - | - | - | 1.954 | [1.0, 4.0] | 386 |
| B_CG | - | - | - | 3 | [3.0, 3.0] | 29347 |
| B_CT | - | - | - | 2.016 | [2.0, 2.0] | 6304 |
| B_GT | - | - | - | 2.638 | [2.0, 3.0] | 165 |
| beta_AC | - | - | - | 2.091 | [1.782, 2.417] | 3515 |
| beta_AG | - | - | - | 1.73 | [1.47, 2.009] | 1166 |
| beta_AT | - | - | - | 0.898 | [0.854, 0.941] | 1646 |
| beta_CG | - | - | - | 1.796 | [1.526, 2.067] | 4819 |
| beta_CT | - | - | - | 1.508 | [1.291, 1.73] | 559 |
| beta_GT | - | - | - | 2.011 | [1.752, 2.283] | 9919 |

**Table S4:**
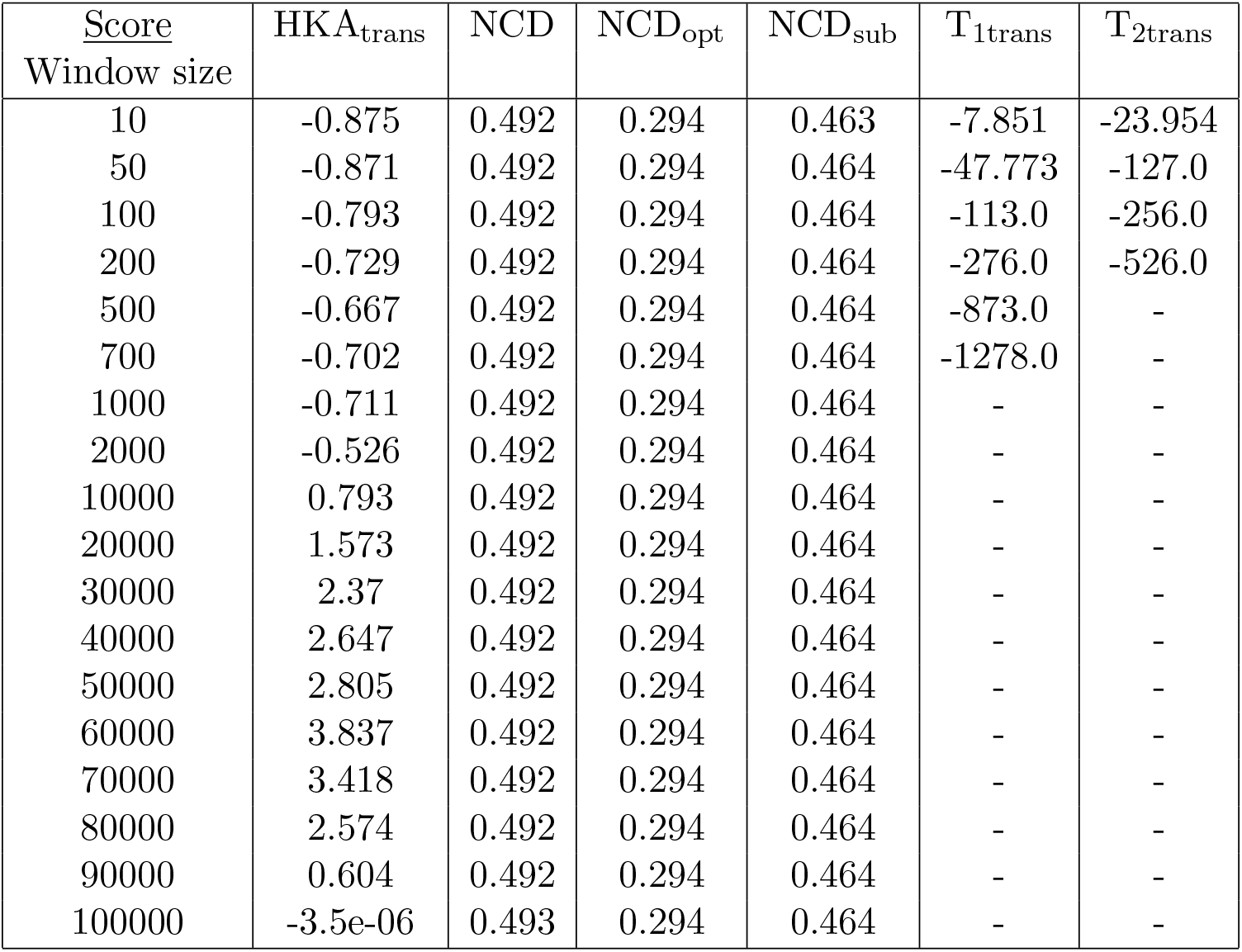
Tests run with MuteBaSS (HKA_trans_, NCD, NCD_opt_, NCD_sub_) and MULLET (T_1trans_, T_2trans_) (Cheng and DeGiorgio, 2019), obtained by averaging the scores with various window sizes and a step size of 10 nucleotides. The data were generated with SLiM on the tree shown in Figure 4 (C) under neutral (drift) conditions.

**Table S5:** Tests run with MuteBaSS (HKA_trans_, NCD, NCD_opt_, NCD_sub_) and MULLET (T_1trans_, T_2trans_) (Cheng and DeGiorgio, 2019), obtained by averaging the scores with various window sizes and a step size of 10 nucleotides. The data were generated with SLiM on the tree shown in Figure 4 (C) under gBGC.

| Score<br>Window size | HKA<sub>trans</sub> | NCD | NCD<sub>opt</sub> | NCD<sub>sub</sub> | T<sub>1trans</sub> | T<sub>2trans</sub> |
| --- | --- | --- | --- | --- | --- | --- |
| 10 | -0.813 | 0.493 | 0.295 | 0.464 | -8.103 | -23.485 |
| 50 | -0.661 | 0.493 | 0.295 | 0.466 | -48.246 | -123.0 |
| 100 | -0.479 | 0.493 | 0.295 | 0.466 | -114.0 | -249.0 |
| 200 | -0.361 | 0.493 | 0.295 | 0.466 | -280.0 | -495.0 |
| 500 | -0.297 | 0.493 | 0.295 | 0.466 | - | - |
| 700 | -0.249 | 0.493 | 0.295 | 0.466 | - | - |
| 1000 | -0.21 | 0.493 | 0.295 | 0.466 | - | - |
| 2000 | -0.127 | 0.493 | 0.295 | 0.466 | - | - |
| 10000 | 0.722 | 0.493 | 0.295 | 0.466 | - | - |
| 20000 | 1.01 | 0.493 | 0.295 | 0.466 | - | - |
| 30000 | 0.52 | 0.493 | 0.295 | 0.466 | - | - |
| 40000 | 0.154 | 0.493 | 0.295 | 0.466 | - | - |
| 50000 | -0.07 | 0.493 | 0.295 | 0.466 | - | - |
| 60000 | -0.77 | 0.494 | 0.295 | 0.466 | - | - |
| 70000 | -0.425 | 0.493 | 0.295 | 0.466 | - | - |
| 80000 | 0.014 | 0.493 | 0.295 | 0.466 | - | - |
| 90000 | 0.042 | 0.493 | 0.295 | 0.466 | - | - |
| 100000 | 0.00072 | 0.493 | 0.295 | 0.466 | - | - |

**Table S6:**
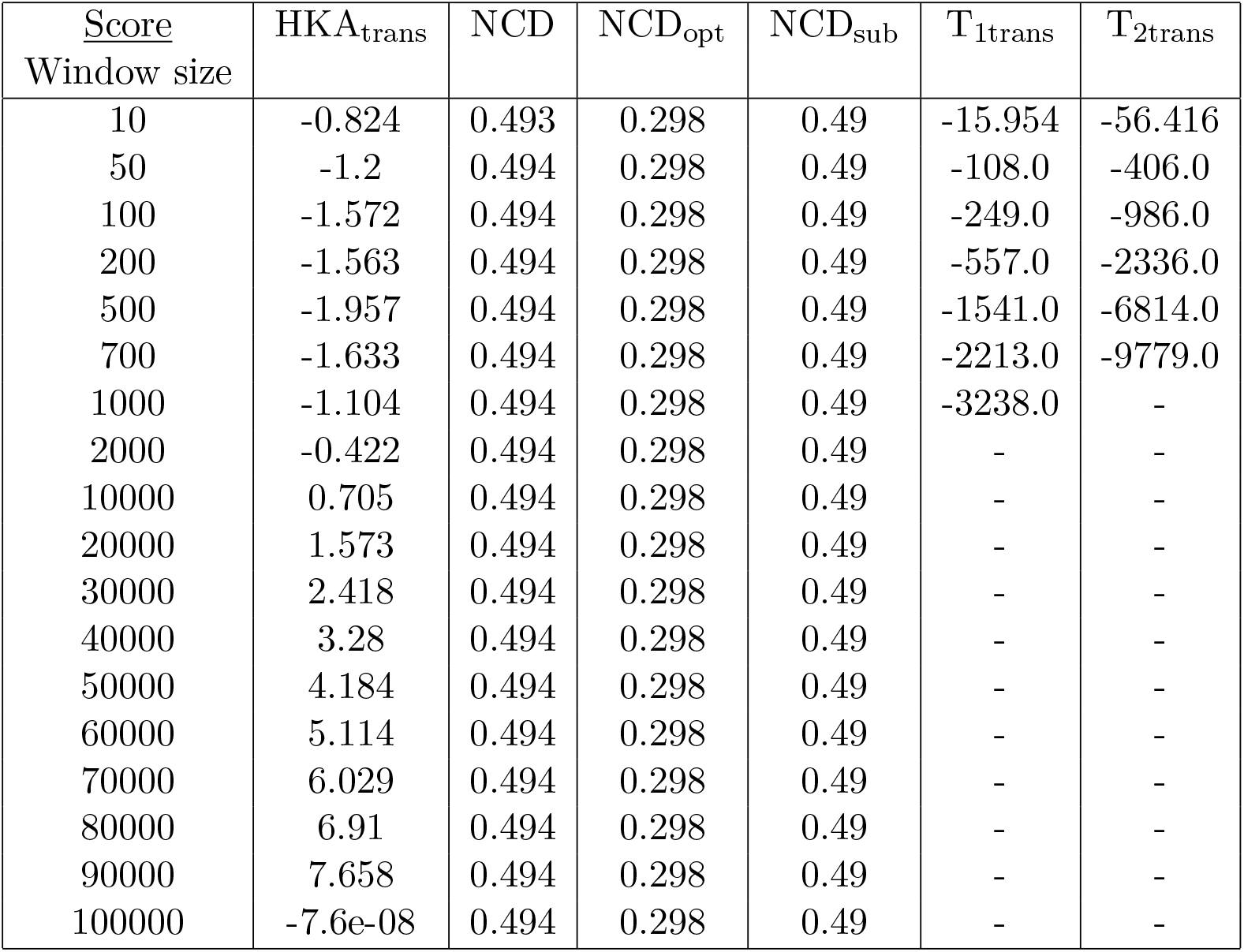
Tests run with MuteBaSS (HKA_trans_, NCD, NCD_opt_, NCD_sub_) and MULLET (T_1trans_, T_2trans_) (Cheng and DeGiorgio, 2019), obtained by averaging the scores with various window sizes and a step size of 10 nucleotides. The data were generated with SLiM on the tree shown in Figure 4 (C) under BS.

| Score<br>Window size | HKA<sub>trans</sub> | NCD | NCD<sub>opt</sub> | NCD<sub>sub</sub> | T<sub>1trans</sub> | T<sub>2trans</sub> |
| --- | --- | --- | --- | --- | --- | --- |
| 10 | -0.824 | 0.493 | 0.298 | 0.49 | -15.954 | -56.416 |
| 50 | -1.2 | 0.494 | 0.298 | 0.49 | -108.0 | -406.0 |
| 100 | -1.572 | 0.494 | 0.298 | 0.49 | -249.0 | -986.0 |
| 200 | -1.563 | 0.494 | 0.298 | 0.49 | -557.0 | -2336.0 |
| 500 | -1.957 | 0.494 | 0.298 | 0.49 | -1541.0 | -6814.0 |
| 700 | -1.633 | 0.494 | 0.298 | 0.49 | -2213.0 | -9779.0 |
| 1000 | -1.104 | 0.494 | 0.298 | 0.49 | -3238.0 | - |
| 2000 | -0.422 | 0.494 | 0.298 | 0.49 | - | - |
| 10000 | 0.705 | 0.494 | 0.298 | 0.49 | - | - |
| 20000 | 1.573 | 0.494 | 0.298 | 0.49 | - | - |
| 30000 | 2.418 | 0.494 | 0.298 | 0.49 | - | - |
| 40000 | 3.28 | 0.494 | 0.298 | 0.49 | - | - |
| 50000 | 4.184 | 0.494 | 0.298 | 0.49 | - | - |
| 60000 | 5.114 | 0.494 | 0.298 | 0.49 | - | - |
| 70000 | 6.029 | 0.494 | 0.298 | 0.49 | - | - |
| 80000 | 6.91 | 0.494 | 0.298 | 0.49 | - | - |
| 90000 | 7.658 | 0.494 | 0.298 | 0.49 | - | - |
| 100000 | -7.6e-08 | 0.494 | 0.298 | 0.49 | - | - |

## References

Andrés, A. M., Hubisz, M. J., Indap, A., Torgerson, D. G., Degenhardt, J. D., Boyko, A. R., Gutenkunst, R. N., White, T. J., Green, E. D., Bustamante, C. D., Clark, A. G., and Nielsen, R. 2009. Targets of Balancing Selection in the Human Genome. Molecular Biology and Evolution, 26(12): 2755–2764.

Bakker, E. G., Toomajian, C., Kreitman, M., and Bergelson, J. 2006. A Genome-Wide Survey of R Gene Polymorphisms in Arabidopsis. The Plant Cell, 18(8): 1803–1818.

Barata, C., Borges, R., and Kosiol, C. 2023. Bait-ER: A Bayesian method to detect targets of selection in Evolve-and-Resequence experiments. Journal of Evolutionary Biology, 36(1): 29–44.

Begun, D. J., Holloway, A. K., Stevens, K., Hillier, L. W., Poh, Y.-P., Hahn, M. W., Nista, P. M., Jones, C. D., Kern, A. D., Dewey, C. N., Pachter, L., Myers, E., and Langley, C. H. 2007. Population Genomics: Whole-Genome Analysis of Polymorphism and Divergence in Drosophila simulans. PLOS Biology, 5(11): e310.

Bitarello, B. D., de Filippo, C., Teixeira, J. C., Schmidt, J. M., Kleinert, P., Meyer, D., and Andrés, A. M. 2018. Signatures of Long-Term Balancing Selection in Human Genomes. Genome Biology and Evolution, 10(3): 939–955.

Bitarello, B. D., Brandt, D. Y. C., Meyer, D., and Andrés, A. M. 2023. Inferring Balancing Selection From Genome-Scale Data. Genome Biology and Evolution, 15(3): evad032.

Borges, R. and Kosiol, C. 2020. Consistency and identifiability of the polymorphism-aware phylogenetic models. Journal of Theoretical Biology, 486: 110074.

Borges, R., Szöllősi, G. J., and Kosiol, C. 2019. Quantifying GC-Biased Gene Conversion in Great Ape Genomes Using Polymorphism-Aware Models. Genetics, 212(4): 1321–1336.

Borges, R., Boussau, B., Szöllősi, G. J., and Kosiol, C. 2022a. Nucleotide Usage Biases Distort Inferences of the Species Tree. Genome Biology and Evolution, 14(1): evab290.

Borges, R., Boussau, B., Höhna, S., Pereira, R. J., and Kosiol, C. 2022b. Polymorphism-aware estimation of species trees and evolutionary forces from genomic sequences with RevBayes. Methods in Ecology and Evolution, 13(11): 2339–2346.

Cagan, A., Theunert, C., Laayouni, H., Santpere, G., Pybus, M., Casals, F., Prüfer, K., Navarro, A., Marques-Bonet, T., Bertranpetit, J., and Andrés, A. M. 2016. Natural Selection in the Great Apes. Molecular Biology and Evolution, 33(12): 3268–3283.

Castric, V. and Vekemans, X. 2004. Plant self-incompatibility in natural populations: A critical assessment of recent theoretical and empirical advances. Molecular Ecology, 13(10): 2873–2889.

Cavalli-Sforza, L. L. and Edwards, A. W. F. 1967. Phylogenetic analysis. Models and estimation procedures. American Journal of Human Genetics, 19(3 Pt 1): 233–257.

Charlesworth, B. and Charlesworth, D. 2010. Elements of Evolutionary Genetics. Roberts and Company.

Charlesworth, D. 2004. Sex determination: Balancing selection in the honey bee. Current biology: CB, 14(14): R568–569.

Charlesworth, D. 2006. Balancing Selection and Its Effects on Sequences in Nearby Genome Regions. PLoS Genetics, 2(4): e64.

Cheng, X. and DeGiorgio, M. 2019. Detection of Shared Balancing Selection in the Absence of Trans-Species Polymorphism. Molecular Biology and Evolution, 36(1): 177–199.

Cheng, X. and DeGiorgio, M. 2020. Flexible Mixture Model Approaches That Accommodate Footprint Size Variability for Robust Detection of Balancing Selection. Molecular Biology and Evolution, 37(11): 3267–3291.

Cheng, X. and DeGiorgio, M. 2022. BalLeRMix+: Mixture model approaches for robust joint identification of both positive selection and long-term balancing selection. Bioinformatics, 38(3): 861–863.

Connallon, T. and Clark, A. G. 2014. Balancing Selection in Species with Separate Sexes: Insights from Fisher’s Geometric Model. Genetics, 197(3): 991–1006.

Croze, M., Wollstein, A., Božičević, V., Živković, D., Stephan, W., and Hutter, S. 2017. A genome-wide scan for genes under balancing selection in Drosophila melanogaster. BMC evolutionary biology, 17(1): 15.

De Maio, N., Schlötterer, C., and Kosiol, C. 2013. Linking Great Apes Genome Evolution across Time Scales Using Polymorphism-Aware Phylogenetic Models. Molecular Biology and Evolution, 30(10): 2249–2262.

De Maio, N., Schrempf, D., and Kosiol, C. 2015. PoMo: An Allele Frequency-Based Approach for Species Tree Estimation. Systematic Biology, 64(6): 1018–1031.

DeGiorgio, M., Lohmueller, K. E., and Nielsen, R. 2014. A Model-Based Approach for Identifying Signatures of Ancient Balancing Selection in Genetic Data. PLOS Genetics, 10(8): e1004561.

Dobzhansky, T. 1955. A review of some fundamental concepts and problems of population genetics. Cold Spring Harbor Symposia on Quantitative Biology, 20: 1–15.

Fernández-Moreno, M. A., Farr, C. L., Kaguni, L. S., and Garesse, R. 2007. Drosophila melanogaster as a Model System to Study Mitochondrial Biology. Methods in molecular biology (Clifton, N.J.), 372: 33–49.

Fijarczyk, A. and Babik, W. 2015. Detecting balancing selection in genomes: Limits and prospects. Molecular Ecology, 24(14): 3529–3545.

Flouri, T., Jiao, X., Rannala, B., and Yang, Z. 2020. A Bayesian Implementation of the Multispecies Coalescent Model with Introgression for Phylogenomic Analysis. Molecular Biology and Evolution, 37(4): 1211–1223.

Haller, B. C. and Messer, P. W. 2019. SLiM 3: Forward Genetic Simulations Beyond the Wright–Fisher Model. Molecular Biology and Evolution, 36(3): 632–637.

Höhna, S., Landis, M. J., Heath, T. A., Boussau, B., Lartillot, N., Moore, B. R., Huelsenbeck, J. P., and Ronquist, F. 2016. RevBayes: Bayesian Phylogenetic Inference Using Graphical Models and an Interactive Model-Specification Language. Systematic Biology, 65(4): 726–736.

Hohna, S., Landis, M. J., and Heath, T. A. 2017. Phylogenetic Inference Using RevBayes. Current Protocols in Bioinformatics, pages 6.16.1–6.16.34.

Höhna, S., Coghill, L. M., Mount, G. G., Thomson, R. C., and Brown, J. M. 2018. P3: Phylogenetic Posterior Prediction in RevBayes. Molecular Biology and Evolution, 35(4): 1028–1034.

Isildak, U., Stella, A., and Fumagalli, M. 2021. Distinguishing between recent balancing selection and incomplete sweep using deep neural networks. Molecular Ecology Resources, 21(8): 2706–2718.

Kelley, J., Walter, L., and Trowsdale, J. 2005. Comparative genomics of major histocompatibility complexes. Immunogenetics, 56(10): 683–695.

Kim, K.-W., Jackson, B. C., Zhang, H., Toews, D. P. L., Taylor, S. A., Greig, E. I., Lovette, I. J., Liu, M. M., Davison, A., Griffith, S. C., Zeng, K., and Burke, T. 2019. Genetics and evidence for balancing selection of a sex-linked colour polymorphism in a songbird. Nature Communications, 10(1): 1852.

Korfmann, K., Gaggiotti, O. E., and Fumagalli, M. 2023. Deep Learning in Population Genetics. Genome Biology and Evolution, 15(2): evad008.

Lanchier, N. 2017. Wright–Fisher and Moran models. In N. Lanchier, editor, Stochastic Modeling, Universitext, pages 203–218. Springer International Publishing, Cham.

Laval, G., Peyrégne, S., Zidane, N., Harmant, C., Renaud, F., Patin, E., Prugnolle, F., and Quintana-Murci, L. 2019. Recent Adaptive Acquisition by African Rainforest Hunter-Gatherers of the Late Pleistocene Sickle-Cell Mutation Suggests Past Differences in Malaria Exposure. American Journal of Human Genetics, 104(3): 553–561.

Lawrence, M. J. 2000. Population Genetics of the Homomorphic Self-incompatibility Polymorphisms in Flowering Plants. Annals of Botany, 85(uppl 1): 221–226.

Mank, J. E. 2017. Population genetics of sexual conflict in the genomic era. Nature Reviews Genetics, 18(12): 721–730.

Moran, P. a. P. 1958. Random processes in genetics. Mathematical Proceedings of the Cambridge Philosophical Society, 54(1): 60–71.

Robinson, M. C., Stone, E. A., and Singh, N. D. 2014. Population Genomic Analysis Reveals No Evidence for GC-Biased Gene Conversion in Drosophila melanogaster. Molecular Biology and Evolution, 31(2): 425–433.

Rozewicki, J., Li, S., Amada, K. M., Standley, D. M., and Katoh, K. 2019. MAFFT-DASH: Integrated protein sequence and structural alignment. Nucleic Acids Research, 47(W1): W5–W10.

Schrempf, D., Minh, B. Q., De Maio, N., von Haeseler, A., and Kosiol, C. 2016. Reversible polymorphism-aware phylogenetic models and their application to tree inference. Journal of Theoretical Biology, 407: 362–370.

Schrempf, D., Minh, B. Q., von Haeseler, A., and Kosiol, C. 2019. Polymorphism-Aware Species Trees with Advanced Mutation Models, Bootstrap, and Rate Heterogeneity. Molecular Biology and Evolution, 36(6): 1294–1301.

Sheehan, S. and Song, Y. S. 2016. Deep Learning for Population Genetic Inference. PLOS Computational Biology, 12(3): e1004845.

Siewert, K. M. and Voight, B. F. 2017. Detecting Long-Term Balancing Selection Using Allele Frequency Correlation. Molecular Biology and Evolution, 34(11): 2996–3005.

Siewert, K. M. and Voight, B. F. 2020. BetaScan2: Standardized Statistics to Detect Balancing Selection Utilizing Substitution Data. Genome Biology and Evolution, 12(2): 3873–3877.

Sprengelmeyer, Q. D., Mansourian, S., Lange, J. D., Matute, D. R., Cooper, B. S., Jirle, E. V., Stensmyr, M. C., and Pool, J. E. 2020. Recurrent Collection of Drosophila melanogaster from Wild African Environments and Genomic Insights into Species History. Molecular Biology and Evolution, 37(3): 627–638.

Spurgin, L. G. and Richardson, D. S. 2010. How pathogens drive genetic diversity: MHC, mechanisms and misunderstandings. Proceedings of the Royal Society B: Biological Sciences, 277(1684): 979–988.

Tajima, F. 1989. Statistical method for testing the neutral mutation hypothesis by DNA polymorphism. Genetics, 123(3): 585–595.

Talts, S., Betancourt, M., Simpson, D., Vehtari, A., and Gelman, A. 2020. Validating Bayesian Inference Algorithms with Simulation-Based Calibration.

Tavare, S. 1986. Some probabilistic and statistical problems in the analysis of DNA sequences. Lectures on Mathematics in the Life Sciences, 17: 57–86.

Yang, Z. 2014. Molecular Evolution: A Statistical Approach. Oxford University Press, Oxford.

Yassin, A., Bastide, H., Chung, H., Veuille, M., David, J. R., and Pool, J. E. 2016. Ancient balancing selection at tan underlies female colour dimorphism in Drosophila erecta. Nature Communications, 7(1): 10400.

Zeng, K., Charlesworth, B., and Hobolth, A. 2021. Studying models of balancing selection using phase-type theory. Genetics, 218(2): iyab055.

